# Architecture of chloroplast TOC-TIC translocon supercomplex

**DOI:** 10.1101/2022.11.20.517165

**Authors:** Hao Liu, Anjie Li, Jean-David Rochaix, Zhenfeng Liu

## Abstract

Chloroplasts rely on the translocon complexes in the outer and inner envelope membranes (termed TOC and TIC, respectively) to import thousands of different nuclear-encoded proteins from the cytosol^1–4^. While previous studies indicated that the TOC and TIC complexes may assemble into larger supercomplexes^5–7^, the overall architectures of the TOC-TIC supercomplexes and the mechanism of preprotein translocation are elusive. Here we report the cryo-electron microscopy (cryo-EM) structure of the TOC-TIC supercomplex from *Chlamydomonas reinhardtii* at an overall resolution of 2.8 Å. The major subunits of the TOC complex (Toc75, Toc90 and Toc34) and TIC complex (Tic214, Tic20, Tic100 and Tic56), three chloroplast translocon-associated proteins (Ctap3, Ctap4 and Ctap5) and three newly-identified small inner-membrane proteins (Simp1-3) have been located in the supercomplex. As the largest protein, Tic214 traverses the inner membrane, the intermembrane space and the outer membrane, connecting the TOC complex with the TIC proteins. An inositol hexaphosphate (InsP6 or I6P) molecule is located at the Tic214-Toc90 interface and stabilizes their assembly. Moreover, four lipid molecules are located within or above an inner-membrane funnel formed by Tic214, Tic20, Simp1 and Ctap5. Furthermore, multiple potential pathways found in the TOC-TIC supercomplex may support translocation of different substrate preproteins into chloroplasts.

---

As the organelles involved in photosynthesis, chloroplasts harbor 2000–3000 nuclear-encoded proteins which are imported from the cytosol through the TOC and TIC complexes^8^. In the past decades, many different components of the TOC and TIC machineries have been identified through biochemical and genetic approaches^4,8^. The TOC complex mainly contains three core subunits named Toc159, Toc34 and Toc75, whereas the TIC complex consists of four major subunits named Tic20, Tic214 (also known as Ycf1), Tic100 and Tic56 (ref.^9^). Besides, several other proteins, such as Tic236, Tic110, Tic62, Tic55, Tic40, Tic32, Tic22 and Tic21, were also reported as components of the TIC complex^2–4^. The TIC complex assembles with the TOC complex to form a large translocon supercomplex with a molecular mass of over 1 megadalton^5–7,10^. Although the protein-import function and components of the TOC-TIC supercomplex have been investigated extensively, little is known about the assembly mechanism of the different subunits of the supercomplex and the translocation pathway for the preproteins remains unclear.

## Overall architecture of the TOC-TIC translocon

As shown in Fig. 1a and b, the TOC-TIC supercomplex from *C. reinhardtii* has an overall shape resembling a torch, and is composed of three complexes, namely TOC, TIC and the intermembrane space complex (ISC) connecting TOC and TIC. TOC mainly contains Toc75, Toc34, Toc90 and a small unidentified chain (chain X) embedded in the outer membrane. Flanking on the Toc90 side, a newly identified β-barrel protein named Ctap4 is located at a closest distance of ~12 Å from TOC (Fig. 1c). For TIC, the transmembrane domains of Tic214 and Tic20 form the core region surrounded by the transmembrane helices of Ctap5, Simp1, Simp2 and Simp3 (Fig.1d). In the intermembrane space, Tic100, Tic56 and the soluble domains of Ctap3, Ctap5, Simp1 and Simp2 intertwine with the intermembrane space domains of Tic214 to form ISC, providing an extended surface to accommodate the intermembrane space domains of Toc90, Toc75, Toc34 and Ctap4. The results of purification, characterization and protein identification of the supercomplex sample are summarized in Extended Data Figure 1 and Supplementary Table 1. The cryo-EM data collection, processing and structure refinement results are presented in Extended Data Figure 2 and Supplementary Table 2.

**Fig. 1:**
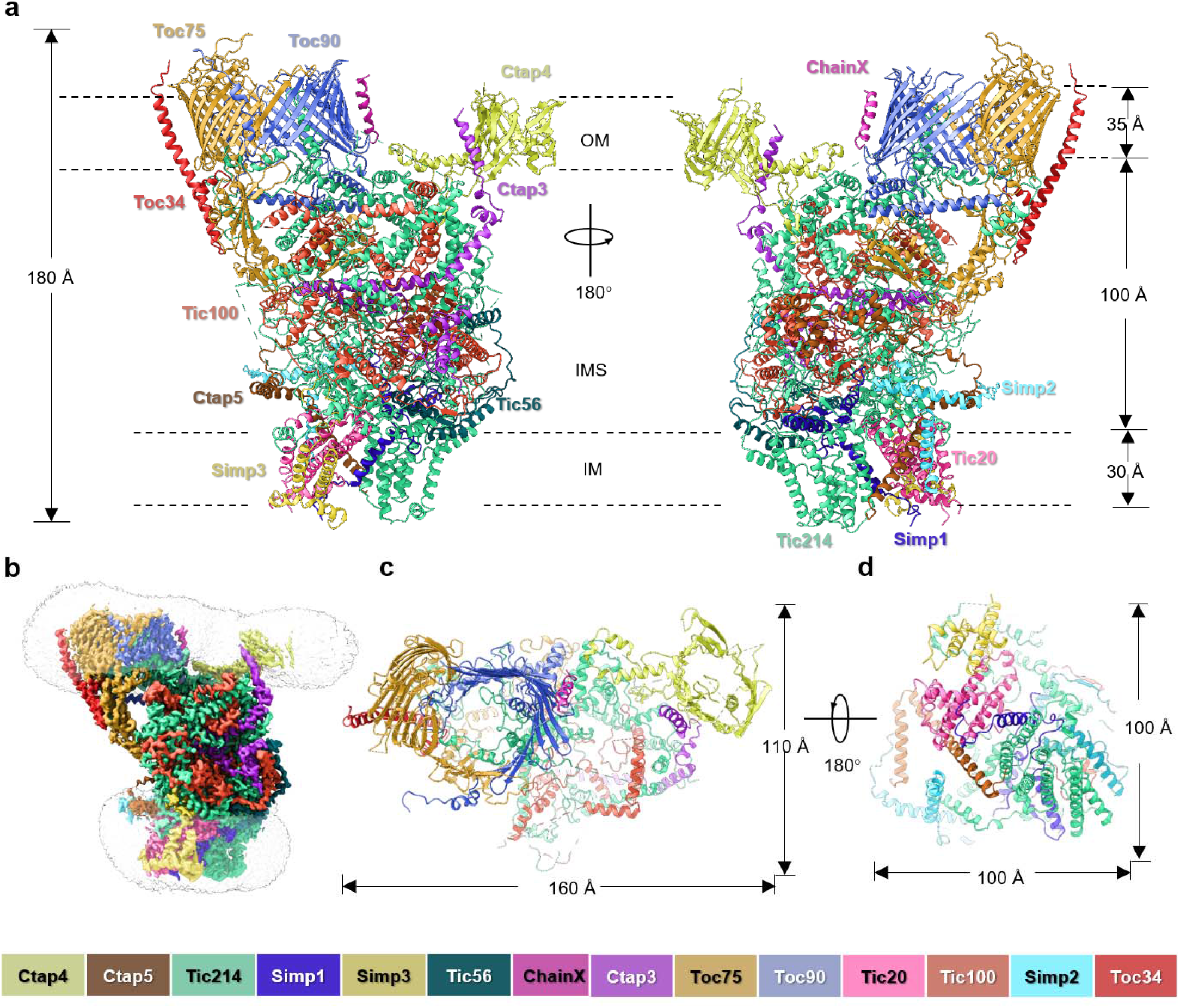
Overall architecture of the TOC-TIC supercomplex. **(a)** Side views of the supercomplex model along the membrane planes. The protein subunits are shown as cartoon models. **(b)** Cryo-EM map of the supercomplex at an overall resolution of 2.77 Å. **(c)** Top view of the supercomplex from the cytosolic side. **(d)** Bottom view of the supercomplex from the chloroplast stromal side.

Curiously, the three-dimensional variability analysis result indicates that the Toc75-Toc90-Toc34 complex and Ctap4 are mobile relative to the ISC and TIC regions (Supplementary Movie 1). They may rotate slightly as rigid bodies by ~3° around a pivot point in the upper side of the ISC region. Ctap4 has a concerted movement along with the Toc75-Toc90-Toc34 complex, as an α-helical domain of Tic214 (Lys602-Ile647, Lys1043-Ile1063 and Gly1454-Gln1491) fills in their gap and bind the two outer-membrane units together.

### The translocon complex in the outer membrane

While previous studies suggested that Toc75 can form a channel by itself^11^ or in complex with Toc159 and Toc34 (ref. ^12^), the cryo-EM structure shows that Toc75 assembles closely with Toc90 at 1:1 ratio to form the channel in the outer membrane (Fig. 2a). Toc34 binds to the peripheral region of Toc75 instead of participating directly in channel formation. The amino-terminal region of Toc75 contains three polypeptide transport-associated domains (POTRA 1–3) which are followed by a membrane-embedded open β-barrel domain with 16 antiparallel β-strands (Fig.2b). POTRA 2 is tilted by ~58° relative to POTRA1, while POTRA2 form a much smaller angle (~35°) with POTRA3. The relationship between POTRA1 and POTRA2 differs largely from the crystal structure of POTRA domains from *Arabidopsis thaliana* Toc75 (ref. ^13^), while the POTRA2–POTRA3 domains of the two structures superpose well (Extended Data Figure 3). In the TOC-TIC supercomplex, POTRA1 is stabilized by nearby proteins (Toc90, Tic214 and Tic100) and forms specific interactions with them, whereas the one in the crystal structure is surrounded by symmetry-related molecules. The transmembrane domain (TMD) of Toc75 curves into a hook-shaped structure (Fig. 2b), for further assembly with the TMD of Toc90 to form an enclosed channel (Fig. 2a, Extended Data Fig. 4).

**Fig. 2:**
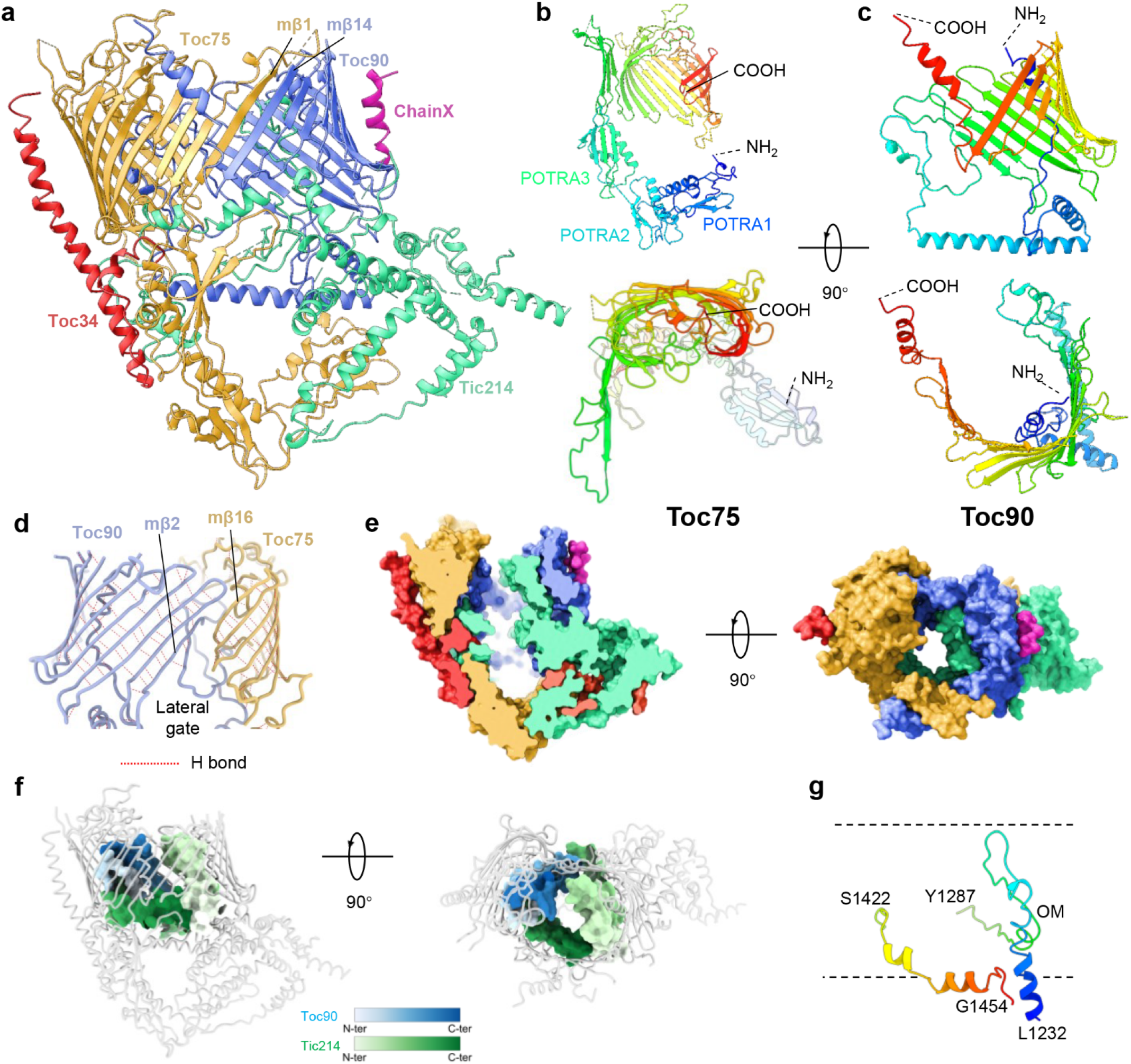
Structure of the Toc75-Toc90-Toc34 complex associated with Tic214. **a,** Cartoon model of the Toc75-Toc90-Toc34-Tic214 complex. **b,** Side view and top view of Toc75 structure colored in rainbow mode. **c**, Side view and top view of Toc90 structure colored in rainbow mode. **d,** A potential lateral gate located between mβ16 of Toc75 and mβ2 of Toc90. The hydrogen bonds (H bonds) between the backbones of adjacent β-strands are indicated by the dotted lines. **e,** Sectional side view and top view of the surface presentation of the Toc75-Toc90-Toc34-Tic214 complex. The color code is the same as the one in a. **f**, The proteins inside the pore lumen are presented as surface models and colored in gradient modes (Toc90, white to blue; Tic214, white to green). **g**, Two motifs of Tic214 inside the channel formed by Toc75 and Toc90. The dash lines indicate the approximate locations of the membrane surfaces.

The TMD of Toc90 contains 14 antiparallel β-strands (Fig. 2c) and forms a C-shaped structure to enclose the adjacent one of Toc75. On one side, the first membrane-embedded β-strand (mβ1) of Toc75 interacts closely with the last membrane-embedded β-strand (mβ14) of Toc90 through backbone hydrogen bonds, forming an antiparallel β-sheet. On the other side, the last membrane-embedded β-strand (mβ16) of Toc75 forms weak van der Waals interactions with the second membrane-embedded β-strand (mβ2, instead of mβ1) of Toc90 (Fig. 2d). Such a weak contact site may potentially serve as a flexible lateral gate, opening for transport of preproteins targeted to the outer membrane, or when the preprotein is too large to fit in the central pore. Toc90 is one of the four members (Toc159, Toc132, Toc120 and Toc90) of the Toc159 family^4^, with a protein-import function similar to that of Toc159 (ref.^14^) and may have a preference for photosynthetic proteins, such as the chlorophyll a/b binding protein (also known as light-harvesting protein/LHCP)^15^. It is evident that the β-barrel domain of Toc90 assembles specifically with Toc75 to form a heterodimeric channel within the outer membrane. The amino-proximal regions of Toc159 and Toc34 contain a GTP-binding domain (G domain) forming a heterodimer or homodimer *in vitro* and *in vivo* ^16,17^. There are some weak cryo-EM densities above the TMDs of the Toc90-Toc75-Toc34 complex, potentially belonging to the G domains of Toc90 and Toc34 (Extended Data Figure 5). The G domains are likely very mobile as the densities are much weaker than the transmembrane domains.

Remarkably, the Toc75-Toc90 channel contains a hollow pore in the middle (Fig. 2e), extending from the cytosolic side to the intermembrane space side. The pore lumen surface is mainly lined by amino acid residues from Toc75 (mβ1-mβ5) and Toc90 (α3-mβ1 loop, mβ1-mβ4, mβ13-mβ14). The α3-mβ1 loop and mβ1-mβ2 loop of Toc90 insert into the cavity of Toc75, and fill the pore lumen of the TOC complex on the Toc75 side (Fig. 2f, blue). Moreover, it is striking that Tic214 also participates in forming the pore lumen surface of the TOC complex by extending two motifs (Leu1232-Tyr1287 and Ser1422-Gly1454) inside the pore (Fig. 2f, green). While the first motif contains a π-shaped helix-loop structure extending from the intermembrane space to the cytoplasmic surface, the second motif harbors two short α-helices located near the exit to the intermembrane space (Fig. 2g). They may contribute to substrate recognition and interaction during translocation of the preproteins through the channel.

Flanking on the Toc90 side, the small β-barrel protein is assigned as Ctap4 (Fig. 1a-c and Extended Data Figure 6). In addition to the β-barrel domain embedded in the outer membrane, the protein contains two hydrophilic motifs (HM1 and HM2) in the intermembrane space region and is associated with the Gln597-Thr663 region of Tic214. Although the local resolution of the β-barrel domain is insufficient for direct identification of the protein, the two hydrophilic motifs are well resolved in the map and exhibit detailed features matching well with the Ctap4 protein (Extended Data Figure 6c). Previously, it was found that seven Ctap proteins (Ctap1-7) including Ctap4 were copurified along with the TOC-TIC supercomplex from *C. reinhardtii*^7^. Our results demonstrate that Ctap4 is a β-barrel membrane protein located in the outer membrane, interacting closely with Tic214 and Ctap3. An α-helix (Val268-Glu283) of Ctap3 is attached to the outer surface of the Ctap4 β-barrel, while the elongated C-terminal domain of Ctap3 is located in the intermembrane space region (Fig. 1a).

### An inositol hexaphosphate molecule mediating Toc90-Tic214 assembly

In an electropositive pocket on the intermembrane-space side of the TOC complex, a small molecule is surrounded by positively-charged residues and sandwiched between Trp1235 of Tic214 and Ser768 of Toc90 (Extended Data Fig. 7a-c). The cryo-EM density matches well with the model of an InsP6 molecule (Extended Data Fig. 7b). The InsP6 molecule is from the endogenous source of *C. reinhardtii* cells and co-purified along with the TOC-TIC supercomplex. The polar extract of small molecules from the purified TOC-TIC supercomplex sample exhibits features similar to those of the InsP6 standard sample in mass spectrometry analysis, confirming that the TOC-TIC supercomplex does contain InsP6 (Extended Fig. 7f and g). The InsP6 closely interacts with seven positively charged residues from Tic214 and Toc90 through salt bridges and four other residues through hydrogen bonds (Extended Fig. 7c-e). Such strong interactions secure the assembly of the pore-lining motif of Tic214 with the inner surface of Toc90 on the intermembrane-space side, as InsP6 wedges in the space between Tic214 and Toc90.

Inositol polyphosphates are phosphorylated derivatives of myo-inositol and have crucial roles in metabolic process regulation, phosphate storage and sensing, hormone signaling and regulation of photosynthetic function in higher plants and algae^18,19^. The discovery of InsP6 in the TOC-TIC supecomplex suggests that InsP6 is crucial for the assembly of the TOC-TIC supercomplex and may also be involved in regulating the process of chloroplast protein transport, as the binding site of InsP6 is at a pivotal interfacial position in the supercomplex.

### The intermembrane-space complex

In the space between the outer and inner membranes, Tic100, Tic214, Tic56, Ctap3, and Ctap5 assemble into a tower-shaped complex measuring ~80–90 Å in height (Fig. 3a). The upper half of intermembrane-space complex (ISC) facing the outer membrane forms a distorted triangular bowl with a shallow cavity in the middle and positive charges enriched at the mouth of the bowl (Fig. 3b), whereas the lower half facing the inner membrane forms a wider quadrilateral base structure (Fig. 3c). On the side surface of the upper half, the ISC provides the binding sites for the POTRA1 and POTRA2 domains of Toc75 as well as the α-helical domains of Toc90 and Ctap4 (Fig. 3a). At the other side near the inner membrane surface, the ISC forms a slightly concaved and flat surface (Fig. 3c) for the association of inner-membrane proteins, such as Tic20, the transmembrane domain of Tic214 and Ctap5 as well as Simp1-3. Thereby, the ISC has a crucial role in connecting TOC with the inner-membrane domains of TIC.

**Fig. 3:**
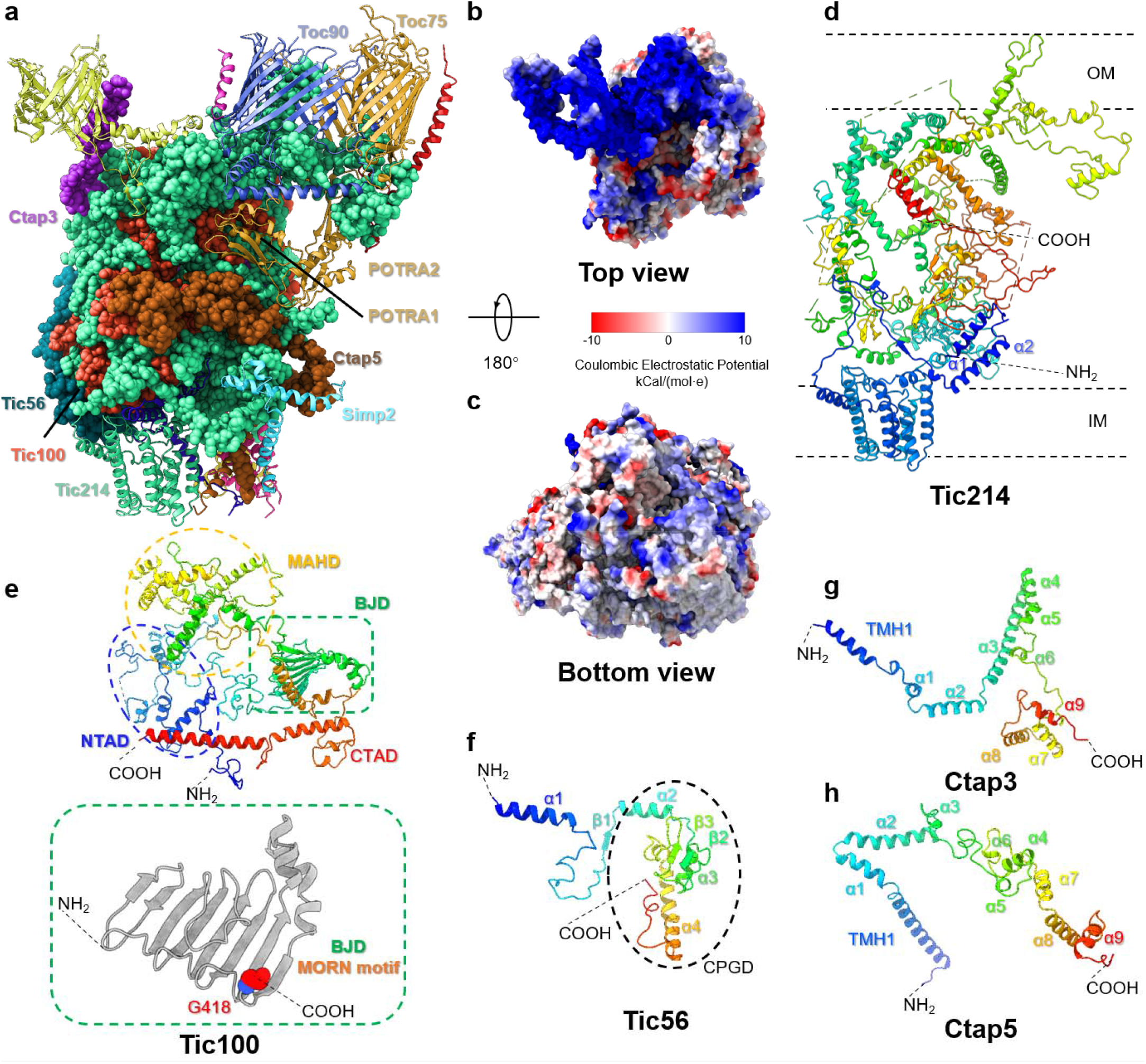
Structural role and components of the intermembrane space complex. **a,** The intermembrane space complex provides binding sites on the surface for the outer-membrane and inner-membrane proteins. The proteins involved in forming the intermembrane space complex are shown as sphere models, while the outer-membrane and inner-membrane proteins are shown as cartoon models. **b** and **c**, Top view and bottom view of the intermembrane space complex from the outer-membrane and inner-membrane sides, respectively. The proteins are presented as surface models, colored by the level of Coulombic Electrostatic Potential (blue, electropositive; white, neutral; red, electronegative). **d-h**, The overall structure of Tic214 (d), Tic100 (e), Tic56 (f), Ctap3 (g) and Ctap5 (h). The proteins are colored in the rainbow mode. Blue, the amino-terminal region; cyan, green and yellow, middle region; red, the carboxy-terminal region. The red sphere in e highlights a glycine residue crucial for the *in vivo* function of Tic100 as reported previously. MORN motif, the Membrane Occupation and Recognition Nexus motif found in Tic100 and potentially involved in binding lipids from the membrane.

Among the five proteins involved in forming ISC, Tic214 is the largest one interacting with nearly all other proteins in the supercomplex. As shown in Fig. 3d, the Tic214 structure mainly consists of ~41 α-helices, two pairs of short antiparallel β-strands and numerous irregular loops. While the amino-proximal region of Tic214 forms a compact transmembrane domain (TMD) embedded in the inner membrane, the middle region folds into an extended and loose structure spanning 70-80 Å across the intermembrane space to connect the TMD with the carboxy-proximal motifs inside the pore lumen of the Toc75-Toc90 channel. The extended structure of Tic214 allows it to connect TOC with TIC proteins and provide a flexible platform for other proteins to bind. The genes encoding Tic214 are essential for the survival of *C. reinhardtii*^20^ and higher plant^21^ cells, and recent biochemical studies indicate that it is a constitutive component of the TOC-TIC supercomplexes^5,7^. Therefore, Tic214 is not only the central subunit for ISC, but is also essential for the assembly of the TOC-TIC supercomplex in general.

Tic100 and Tic56 are both soluble proteins important for the assembly of the TIC complex and essential for accumulation of photosynthetic proteins in plants^5^. The structure of Tic100 has an overall shape of a wreath with four major domains, namely the amino-terminal α-helical domain (NTAD), β-jellyroll domain (BJD), middle α-helical domain (MAHD) and the carboxy-terminal α-helical domain (CTAD) (Fig. 3e). These domains are mainly connected through flexible loops and only a few non-covalent interactions are observed between NTAD and MAHD or between BJD and CTAD. Instead of forming a compact structure by itself, Tic100 intertwines with the intermembrane-space domain of Tic214 to form the central part of ISC. The Arabidopsis *tic100* mutant has reduced level of 1 MDa Tic complex and lower chloroplast protein import rate in comparison with the wild type, and mutation of a glycine residue (corresponding to Gly418 in *Cr*Tic100) located at the BJD domain (Fig. 3e) to Arg led to partial loss of function of Tic100^22^. Therefore, Tic100 is crucial for the assembly of the TIC complex by forming close interactions with Tic214 and may facilitate the process of protein import into chloroplasts by establishing a physical bridge between the TOC and TIC complexes.

Tic56 is a smaller protein with a short α-helix (Gly78-Thr93) at the N-terminal region connected with a carboxy-proximal globular domain (CPGD, Pro127-Arg244) through a long loop (Fig. 3f). While the amino-terminal α-helix of Tic56 interacts with the carboxy-terminal α-helix of Tic100, the CPGD of Tic56 intercalates at the cleft between the transmembrane domain of Tic214 and CTAD of Tic100 to stabilize their assembly (Fig. 3a). The absence of Tic56 in *Arabidopsis* also leads to reduction of the 1 MDa Tic complex and the chloroplasts are deficient in protein import^5,23^.

Ctap3 has an extended α-helical domain (Arg295-Ser472) at the carboxy-proximal region (Fig. 3g). It docks two short α-helices (α1 and α2) on the outer surface of the upper half of ISC, inserts a long α-helix (α3) and two short ones (α4 and α5) into the bottom of the bowl-like structure formed by Tic214 and Tic100 (Fig. 3b). The last three α-helices (α7-9) of Ctap3 bind to the amino-terminal helix of Tic56 and the carboxy-terminal helix of Tic100. On the opposite side, Ctap5 also has an extended carboxy-proximal α-helical domain (Glu215-Ala329, Fig. 3h) attached on a lower groove surface of the Tic100-Tic214 complex below the POTRA1 and POTRA2 domains of Toc75 (Fig. 3a). It interacts closely with the BJD of Tic100 and Tic214 (Gln1621-Phe1645 and Tyr1917-Ile1949) and provides binding sites for the POTRA1 and POTRA2 domains of Toc75. Thereby, Ctap5 stabilizes the Tic214-Tic100 complex and serves as a bridge to connect Toc75 with the inner-membrane complex.

### The translocon complex in the inner membrane

Tic20 is a crucial component of the TIC complex, interacting closely with a preprotein substrate during the late stage of the import process^24^. Mutant plants with reduced Tic20 levels exhibit inefficient import and accumulation of plastid preproteins, reduced photosynthetic capacity and growth defects^25,26^. As shown in Fig. 4a and b, Tic20 assembles with the transmembrane domain of Tic214 and four small membrane proteins (Ctap5, Simp1, Simp2 and Simp3) to form the translocon complex embedded in the inner membrane. There are 15 transmembrane helices (TMH) in TIC, arranged in an asymmetric way within the membrane. Tic20 and Tic214 contain four and six TMHs respectively, while the remaining five are from Ctap5, Simp1, Simp2 and Simp3 (Fig. 4c-g, Extended Data Figure 8). At the peripheral region of Tic20, Simp3 associates with TMH3 and α3 of Tic20 to stabilize the local structure around Tic20 (Fig. 4b). Besides, Simp3 also interacts closely with Tic214. The carboxy-terminal tail of Simp3 attaches to a surface pocket surrounded by the three parts (Pro468-Ile475, Arg735-Lys749 and Lys815-Tyr825) of Tic214, and a short helix-loop-helix motif (Arg426-Lys447) of Tic214 connects Simp3 (TMH2) with Tic20 (TMH3-α2). Remarkably, TMH1 and TMH4 from Tic20, TMH1 from Simp1, TMH3 and TMH6 from Tic214 and TMH1 from Ctap5 collectively form a funnel-like structure in the middle region, resembling the SecY protein-conducting channel^27^ (Fig. 4h and i). On the intermembrane-space side, the funnel is covered by an α-helix(α6)-loop motif (Gly328-Val380) following TMH6 of Tic214 and the TMH1-α1b loop region of Tic20 (Fig. 4a). Instead of having a substrate protein inside the funnel, three lipid molecules are located inside the pore and a fourth one is found in a pocket above the funnel (Fig. 4j, Extended Data Fig. 9a). The lipid molecules form hydrogen bonds and hydrophobic interactions with amino acid residues from Tic20, Ctap5, Tic214 and Simp1 (Extended Data Fig. 9b, c). At the bottom side facing the stroma, a phospholipid molecule (PG604) serves as a plug to occlude the bottleneck and prevent leakage of the funnel (Fig. 4j). Besides, the funnel has a lateral gate opened toward the lipid bilayer (Fig. 4h). A similar lateral gate was also found in SecY, serving as an exit for the hydrophobic signal peptide segment and/or the membrane-spanning domains of substrate proteins (Fig. 4i)^27^. Moreover, TMH1 of Simp2 is located at the edge of the lateral gate, defining the peripheral boundary of the exit gate (Fig. 4b). Additionally, numerous other lipid (and detergent) molecules are located at the peripheral region of the TIC complex (Extended Data Fig. 9d).

**Fig. 4:**
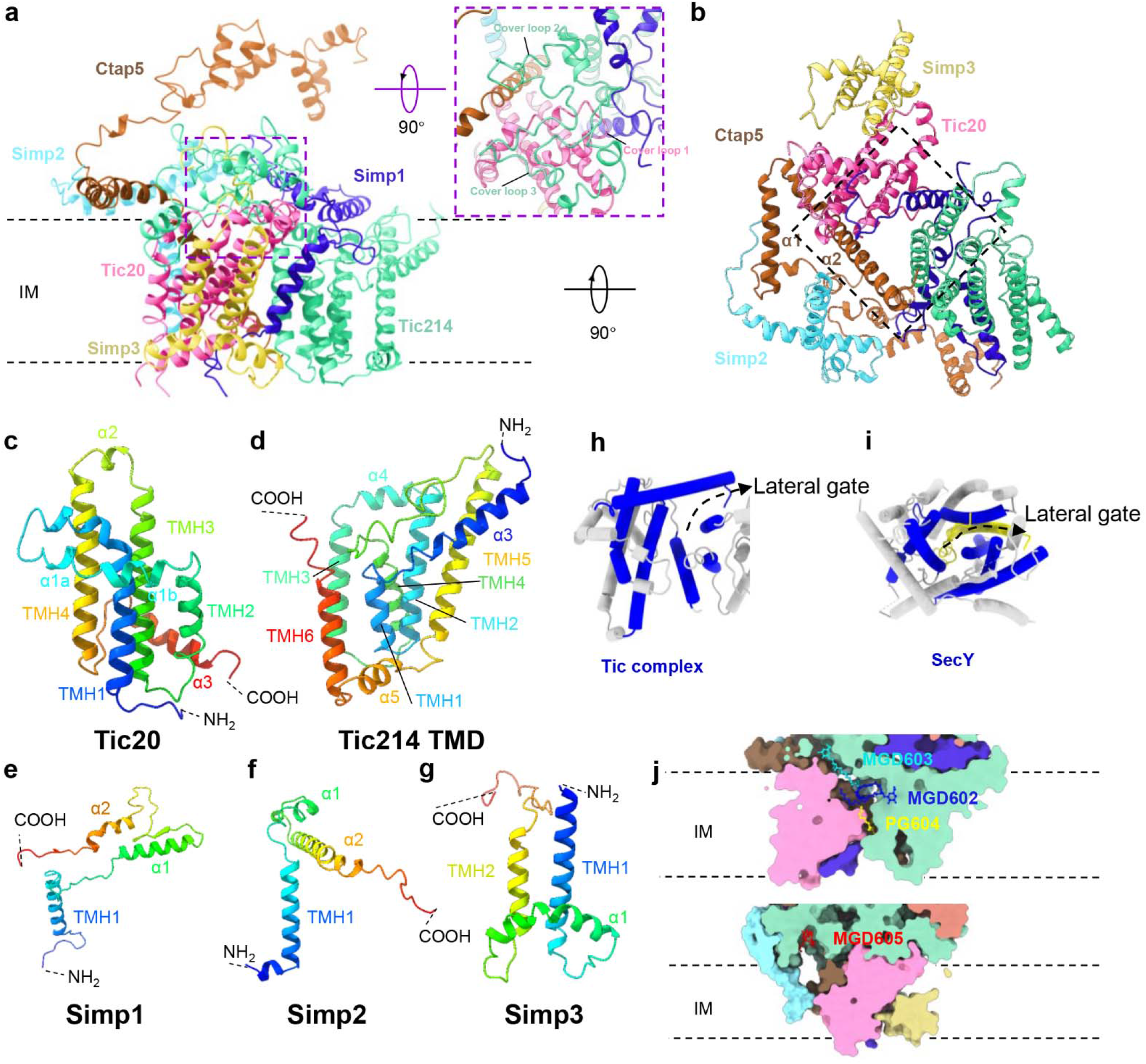
Arrangement of protein subunits and lipid molecules in the inner membrane complex. **a and b,** side view and bottom view of the translocon complex in the inner membrane. In a, the area in the purple dashed box contains the loops from Tic20 and Tic214 covering the funnel entrance underneath. The view in b is from the stromal side and the area in the dark dashed box contains a central funnel filled with lipid molecules (omitted for clarity). **c-g,** structures of Tic20 (c), Tic214 transmembrane domain (TMD, d), Simp1 (e), Simp2 (f) and Simp3 (g). **h**, the central funnel-like region of Tic20-Tic214-Ctap5-Simp1 complex. The view is from the intermembrane space side. **i**, The protein-conducting channel of SecY with substrate bound. The transmembrane helices encircling the central pore are colored in blue, while the other parts are in silver. The polypeptide substrate of SecY is in yellow. (PDB code: 5EUL). **j**, Binding sites of four lipid molecules at the subunit interfaces. MGD, monogalactosyl-diacylglycerol; PG, phosphatidyl glycerol.

### Putative protein translocation pathways

TOC contains a straight open pore running from the cytosolic side to the intermembrane space (Fig. 5a). The pore appears to have a bottleneck near the cytoplasmic entrance, measuring 11.7 Å wide in the shortest dimension and 20.5 Å wide in the longest dimension (Fig. 5b). It is surrounded by amino acid residues from the mβ1-mβ2 loop region of Toc90 as well as the mβ3-mβ4 loop and mβ7-mβ8 of Toc75 (Fig. 5b). The surface of the pore lumen is electropositive on one side and electronegative on the other side (Extended Data Fig. 10a and b). While the entrance appears wide enough to accommodate an extended polypeptide of the substrate preprotein, the flexible loops around the entrance may adjust dynamically according to different parts of the translocated preprotein. Below the entrance, the pore diameter increases to 14–22 Å, wider than the entrance (Fig. 5b and c). There are two exits on different sides of the bottom. While one exit is open directly to the intermembrane space, the other one is connected with a slide-like groove formed by POTRA2–POTRA3 of Toc75 and the nearby parts of Tic100, Tic214 and Ctap3 (Extended Data Fig. 10c and d). Such a groove may help to guide the preproteins to the cavity above the central funnel of the TIC complex or to the adjacent portal through the triangular entrance by Ctap5.

**Fig. 5:**
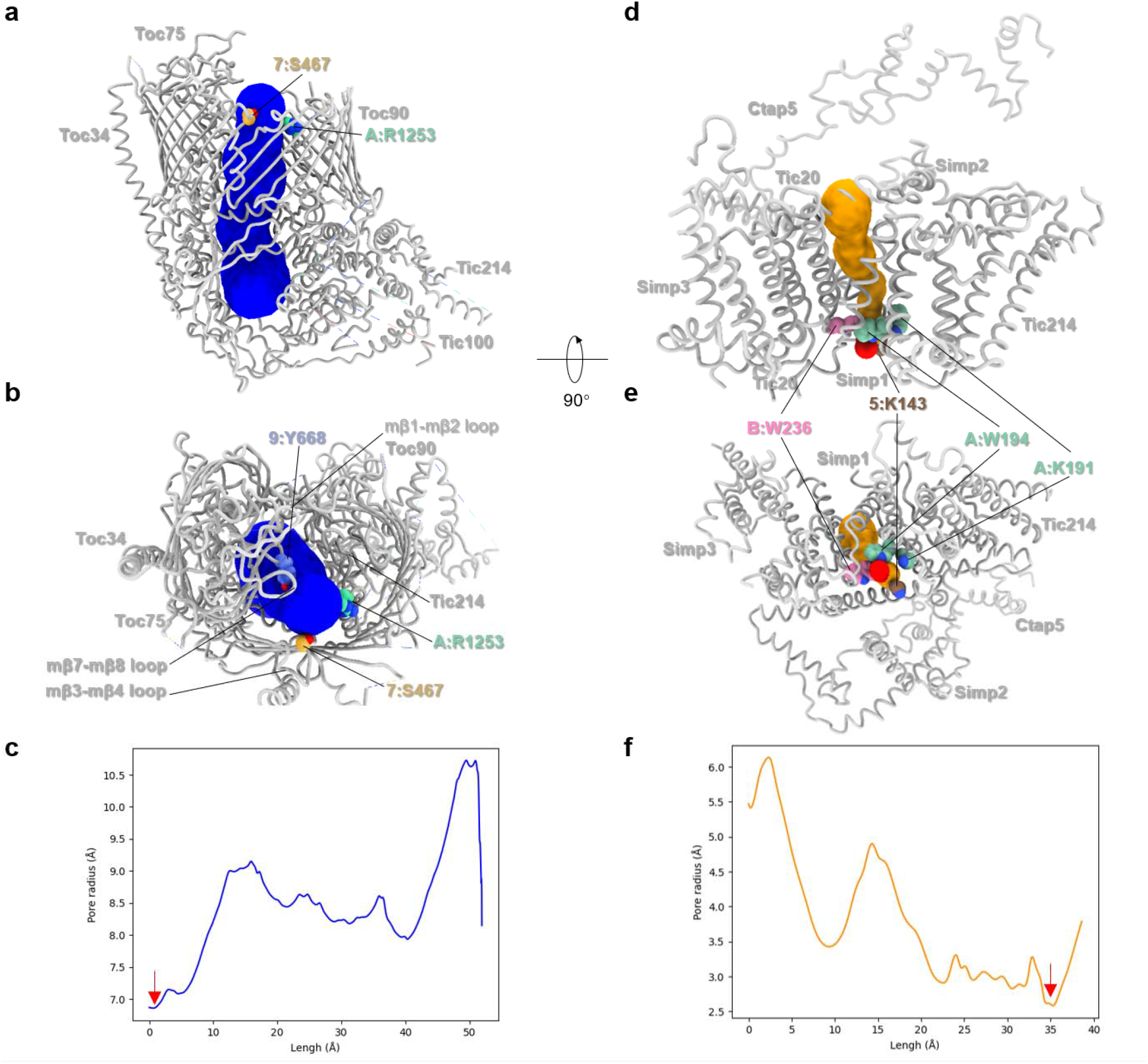
The intrinsic pores found in the TOC and TIC complexes. **a and b**, The presence of a continuous wide pore running through the TOC complex. The putative pore is presented as a transparent blue tube model. Side view (a) and top view (from cytoplasmic side, b) of the TOC complex are shown. Three key amino acid residues at the cytoplasmic entrance are shown in sphere models and the labels indicate chain name:residue name and number. 7, Toc75; A, Tic214; 9, Toc90. **c**, Distribution of pore radius along the central axis of the TOC pore. The red arrow indicates the constriction site of the pore with the smallest pore radius. **d** and **e,** The central pore of the TIC complex shown as a transparent orange tube model. The region in the red is the constriction site. Side view (d) and bottom view (from stromal side, e) of the TIC complex are shown. Four key amino acid residues at the constriction site near the stromal surface are presented in sphere models and labeled by chain name:residue name and number. A, Tic214; B, Tic20; 5, Ctap5. **f**, Distribution of pore radius along the central axis of the TIC pore. The red arrow indicates the constriction site of the pore with the smallest pore radius.

The pore in the central funnel between Tic214 and Tic20 is overall much narrower than the one in the TOC complex (Fig. 5d), indicating that the TIC channel may be in a closed state. The upper half of the pore is 7.0–12.1 Å wide, while the lower half is only 5.2–6.6 Å wide (Fig. 5e and f). The constriction site is located near the exit at the stromal surface and surrounded by amino acid residues from Tic214, Ctap5 and Tic20 (Fig. 5e). Therefore, the TIC pore will need to expand by adjusting local protein conformations dramatically for the preprotein to enter and go through. As the transmembrane helices of Ctap5 and Simp1 are not tightly restrained by nearby subunits, they might be able to move outward and adjust flexibly in response to the preprotein being translocated.

The TOC and TIC complexes are responsible for translocation of various nucleus-encoded proteins into different chloroplast subcompartments^1,3^. As different substrate preproteins have distinct surface properties and target locations, they might be translocated through different pathways. For soluble proteins targeted to the stroma, such as the small subunit of ribulose 1,5-bisphosphate carboxylase (RBCS) or ferredoxin (Fd)^10,24^, they will need to pass through the central pore of the Toc75-Toc90 channel firstly. Guided by the surface groove on ISC, they may efficiently traverse the intermembrane space and are further translocated across the inner membrane, presumably through the central funnel within the Tic20-Tic214-Ctap5-Simp1 complex (Fig. 6a, Exit 1). Because the central funnel of the Tic20-Tic214-Ctap5-Simp1 complex is covered by the flexible loops from Tic214 and Tic20 (Fig. 4a), opening of the entrance will require rearrangement of the cover loops for the preproteins to enter. Subsequently, the central pore needs to expand to accommodate the polypeptides of the preproteins, so that they can pass through the pore and reach the stroma. For the preproteins targeted to the intermembrane space, such as the Tic22 protein^28^, they may exit the Toc75–Toc90 channel through a side portal (Exit 2) outlined by the POTRA1–POTRA2 of Toc75 and α3 of Toc90, or through the adjacent one (Exit 3) between POTRA2–POTRA3 of Toc75 and the bowl-like structure of the Tic214–Tic100 complex (Fig. 6b). For preproteins targeted to the inner membrane^29^, there are two potential lateral portals allowing them to be translocated into the membrane, after they pass through the Toc75-Toc90 channel (Fig. 6c). One lateral portal (exit 4) is connected with the central funnel of Tic20-Tic214-Ctap5-Simp1 and the other one (exit 5) is on the adjacent side of Simp2 along the surface of TM1-TM2 of Tic20. For the preproteins to be released through exit 4, they may need to enter the portal through a triangle-shaped entrance outlined by two amphipathic helices (α1 and α2) of Ctap5 above the portal (Extended Data Fig. 10d, e). The lateral portals may also support translocation of photosynthetic membrane proteins, such as LHCP and others, by providing transitional docking sites for their transmembrane domains.

**Fig. 6:**
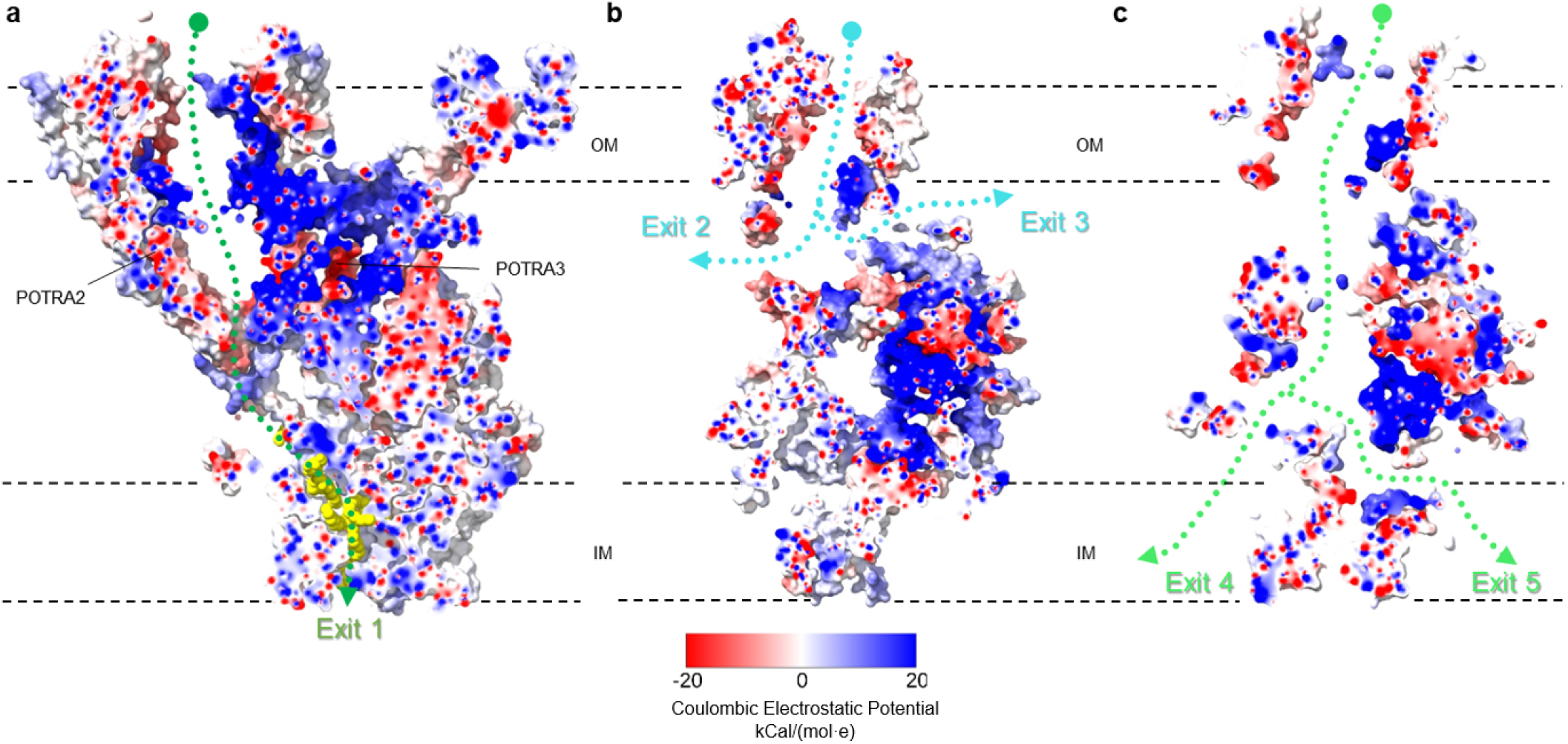
The potential preprotein translocation pathways in the TOC-TIC supercomplex. **a-c** The potential translocation pathways for soluble proteins or LHCP proteins targeted to stroma (a), proteins targeted to the intermembrane space (b) or membrane proteins targeted to the inner membrane (c). The green and cyan dotted lines indicate the approximate locations of the translocation pathways, while the arrows show the possible exit portals for different substrate preproteins. The TOC and TIC proteins are presented as surface models colored by the level of Coulombic Electrostatic Potential (blue, electropositive; white, neutral; red electronegative), whereas lipid molecules are shown as yellow spheres. The sectional views of the TOC-TIC supercomplex oriented in three different angles are shown in a-c.

## Discussion

The cryo-EM map reveals a TOC complex from *C. reinhardtii* composed of Toc90, Toc75 and Toc34 with a stoichiometry of 1:1:1, whereas previous studies on plant TOC complexes suggested that the stoichiometry of Toc159:Toc75:Toc34 is 1:4:4-5 or 1:3:3 (ref.^12,30^). Thus, much larger TOC complexes may exist in plants. Alternatively, the Toc159-Toc75-Toc34 complexes with 1:1:1 ratio might coexist with the Toc75-Toc34 complexes with no Toc159 bound, accounting for the excess molar ratio of Toc75 and Toc34 relative to Toc159. Previously, it was reported that Toc75 or Tic20 alone can form channels by itself when reconstituted in liposomes^31,32^, whereas our work demonstrates that Toc75 assembles with Toc90 to form the TOC channel and is further connected with the TIC complex formed by Tic20, Tic214, Simp1, Simp2, Simp3 and Ctap5. Nevertheless, it can not be ruled out that Toc75 or Tic20 may still form channels alone, in case of free Toc75 or Tic20 proteins in the membrane.

It was reported that Tic236 serves as a large protein linking the TOC and TIC complexes^33^, whereas no density corresponding to Tic236 can be found in the map. Tic214 and Tic100, instead of Tic236, form the ISC core to link the TOC and TIC complexes (Fig. 3a). Nevertheless, previous work suggested that Tic110 may form a channel sensitive to transit peptides and function as a translocation pore for the preproteins at the inner membrane^34^. Another biochemical study on plant TIC complex indicated that Tic21 is loosely associated with the complex^10^. In the cryo-EM map of the TOC-TIC supercomplex from *C. reinhardtii*, neither Tic110 nor Tic21 is observed, suggesting they are either lost during purification or may exist in very low abundance. Alternatively, Tic110 may function downstream of the translocon channel instead of being part of the TOC-TIC supercomplex^8^.

When a tightly folded protein is used as the substrate for the TOC-TIC translocon, the pore may open to a size greater than 25.6 Å^35^, much larger than the pores observed in the structure (Fig. 5). In that case, the TOC and TIC complexes will need to go through large conformational changes in the pore-lining regions to accommodate the folded proteins. Recently, a Ycf2-FtsHi heteromeric AAA-ATPase complex was identified to be the import motor associated with the TIC complex from *A. thaliana*, utilizing the energy from ATP to drive the import process^36^. In *C. reinhardtii*, three FtsH-like AAA-proteins associated with Tic20 and Tic214 were also identified^7^. In the TIC complex, there is a large concaved groove between the TMD of Tic214 and Simp3 (Extended Data Figure 10f). Potentially, the groove and the U-shaped stromal surface of the TIC complex may provide binding sites for the import motor. In the cryo-EM map, no extra densities corresponding to the FtsH-like AAA proteins are visible, suggesting that they might also be lost during the purification process or their association with the TIC complex relies on the presence of preprotein substrates. Therefore, the TOC-TIC supercomplex structure reported hereby represents the pre-import state without any preprotein substrates or AAA-ATPase bound.

## Methods

### Purification and biochemical analysis of the TOC-TIC supercomplex

The *C. reinhardtii* CS1_FC1D12 strain^37^ from the *Chlamydomonas* Resource Center expressing Tic20-Venus-3×Flag fusion protein was used for purification of the TOC-TIC supercomplex through affinity^7^ and size-exclusion chromatography methods.

For protein expression, the single colony selected with 25 μg/ml paromomycin was used to inoculate 100 mL Tris-acetate-phosphate medium with 4 × Kropat’s Trace Elements^38^ . After the cell density reached an OD_600nm_ of 1.0, the culture was then used to inoculate 2 l TAP medium with constant bubbling of filtered air through pumps at 27 °C under 150 μmol m^−2^ s^−1^ photons light conditions for 7 days. The cells were harvested through centrifugation at 5,053 ×g in JLA-8.1 rotor (Beckman) and the pellets were stored at −80 °C.

Throughout the protein purification process, all steps were carried out at 4 °C. For each preparation, 50 g frozen cell pellets were resuspended in a lysis buffer [50 mM HEPES-KOH pH 7.5, 250 mM NaCl, 10% (v:v) Glycercol, 5 mM EDTA, 4 mM DTT, 1×Protease Inhibitor Cocktail (EDTA-Free,100 × in DMSO, MedChemExpress/MCE)] at 1:10 (m:v) ratio, yielding a suspension with about 1 mg/mL chlorophyll (Chl)., The suspension was homogenized by using the T10 basic homogenizer (IKA). The cells were lysed by passing through a high-pressure homogenizer (ATS Engineering, Shanghai) 5 times at 1,000 bar. Lauryl maltose neopentyl glycol (LMNG, NG310, Anatrace) and Digitonin (D-3203, BIOSYNTH) were added to the cell lysate at 1.0% (w/v) and 0.2% (w/v) final concentration respectively, to solubilize the membrane protein complexes. 1 mL MonoRab™ Anti-DYKDDDDK Magnetic Beads (L00835, GenScipt) was added to the mixture to capture the target protein. After the mixture was stirred for 2 h, the magnetic beads were isolated and washed at least 5 times by using the wash buffer [50 mM HEPES-KOH pH 7.5, 150 mM NaCl, 10% (v:v) Glycercol, 1mM DTT, 0.06% Digitonin]. The target protein complexes were eluted twice with 500 μg/ml 3× DYKDDDDK peptide (RP21087, GenScript) dissolved in the wash buffer. The eluent was concentrated to ~0.5 ml and then applied to the size-exclusion chromatography (SEC) column (Superose 6, 10/300, GE Healthcare) in the SEC buffer (50 mM Tris-HCl, pH 7.5, 150 mM NaCl, 0.06% Digitonin). The fractions containing the TOC-TIC supercomplex were pooled and concentrated to 16 μl in a 100-kDa MWCO Amicon Ultra-0.5 Centrifugal Filter Unit (UFC5100, Millipore).

### Protein identification

For protein identification, the TOC-TIC supercomplex sample was analyzed firstly through sodium dodecyl sulfate-polyacrylamide gel electrophoresis (SDS-PAGE, using ExpressPlus™ PAGE Gel, 10×8, 4-12%, GenScipt), or through the Blue Native-polyacrylamide gel electrophoresis (BN-PAGE)^39^. The gels were fixed and stained by using the Silver Stain for Mass Spectrometry kit (C500021, Sangon Biotech). Each individual protein band from the SDS-PAGE or BN-PAGE was excised and silver was removed before the mass spectroscopy analysis.

For preparing the peptide sample from the protein bands of SDS-PAGE, the protein samples were treated with DTT and iodoacetamide, and trypsin was added to the samples to digest the proteins overnight. Subsequently, the products were desalted by using C18 ZipTip column, mixed with the Alpha-cyano-4-hydroxycinnamic acid (CHCA), and then spotted on the plate. The data were acquired through the matrix assisted laser desorption/ionization time-of-flight/time-of-flight mass spectrometer (MALDI-TOF/TOF UltraFlexTreme^™^, Brucker, Germany). Database search was performed by using the Peptide Mass Fingerprint or MS/MS ION Search page from a local MASCOT website. Cysteine carbamidomethylation was set as a fixed modification, while methionine oxidation was set as a variable modification. The SWISSPROT database was used to establish a reference database for MALDI-TOF MS spectra.

For peptide samples prepared from the BN-PAGE bands, the samples were analyzed by using the nano LC-Q EXACTIVE system (Thermo Scientific). The protein samples were treated with DTT and iodoacetamide, and then trypsin enzymatic solution was added. The mixtures were incubated for 30 min, and then chymotrypsin was added to digest the proteins overnight. Afterwards, the peptides were separated by passing the samples through the C18 reversed-phase column (7.5 μm × 20 cm, 3 μm) at a flow rate of 300 nl/min. The mobile phase consisted of two components, namely phase A (0.1% Formic acid aqueous) and phase B (0.1% Formic acid Acetonitrile). The column was first eluted with a linear gradient from 4% to 8%B from 0 to 8 min and then with a linear gradient from 22% to 32%B from 8 min to 58 min, followed by an equilibration step for 7 min with 90%B. The MS analysis was performed with a scan range of 300-1600 m/z for MS, ion spray voltage was set at 2.0 kV in the positive mode, and the micropipette temperature was 320 ℃. The data collected were used for searching against the UniProt_Proteome_Chlamydomonas reinhardtii_2021 database to find matching fragments by using the Thermo Proteome Discoverer program (1.4.0.288).

### Cryo-EM sample preparation and data collection

For cryo-EM sample preparation, 3.0 μl of the supercomplex sample was added onto a H_2_/O_2_ glow-discharged (in Solarus 950, Gatan) holey carbon grid coated with 2 nm ultrathin carbon film (Quantifoil 300-mesh, R 2/2). After waiting for 60 s, the grids were blotted for 8 s with a force level of 0 and immersed into liquid ethane using a Vitrobot (Mark IV, Thermo Fisher Scientific) under 100% humidity at 4 °C. The frozen grids were then transferred to liquid nitrogen for storage.

The images were collected by using a system comprised of a Titan Krios G3 (Thermo Fisher Scientific) operating at 300 kV and a K2 Summit detector (Gatan). Movie stacks were automatically acquired in the super-resolution mode (with super-resolution pixel size 0.675 Å) using the SerialEM program, with a defocus range from −1.5 μm to −1.8 μm. The total dose was approximately 50 e^−^/Å^2^ in 33 frames. The images were recorded through the beam-image shift data collection method^40^.

### Cryo-EM data processing and refinement

The workflow for cryo-EM data processing is summarized in Extended Data Figure 2. The procedures described below were performed by using cryoSPARC v3.2 (ref. ^41^) unless stated otherwise. Firstly, 9,984 movie stacks were motion-corrected by Patch-based motion correction and dose-weighted with two-fold binning. Micrograph contrast transfer functions were estimated by Patch-based CTF estimation.

For particle picking, an initial round was performed using manual picking and the Blob Picker Tuner. After the first round of 2D classification, ab-initio reconstruction and Non-Uniform refinement^42^ were carried out. The 3D map exhibited a severe preferred orientation due to lack of top-view and bottom-view particles. To pick the particles with more orientations, simulated data were used to generate particles with all orientations. Subsequently, a 2D classification step using the simulated data was used to generate templates for the Template picking process. Around 6,814,089 particles were picked by using Template picking and extracted with four-fold binning.

For 3D classification in Relion 4.0 (ref. ^43^), all particles were converted to star format by using cs2star 0.4.1. 3D classification (Classes=6, T=6) was carried out to generate 6 reference maps by using the refined model obtained in the last non-uniform refinement step as a reference. All the maps were imported to cryoSPARC, and the corresponding selected particles were re-extracted with two-fold binning. Following a heterogeneous refinement and a non-uniform refinement job, a reconstruction at an overall resolution of 2.9 Å was obtained. Then, two rounds of heterogeneous refinement followed by Local CTF Refinement, Global CTF Refinement, Remove Duplicate Particles, Local motion correction and non-uniform refinement were performed and further improved the overall map resolution to 2.77 Å.

As the region around the TOC complex has much weaker densities than the TIC complex region, a mask covering the TOC complex and nearby outer membrane domains was generated by using UCSF ChimeraX 1.4 (ref. ^44^) and Volume Tools in cryoSPARC, and applied to improve the local map quality. A combined process with Local Refinement and Global CTF Refine was carried out in two rounds to generate a local map at 3.40 Å resolution. After the refinement, the map quality around the TOC complex in the local map appears much better than the corresponding region in the global map.

The cryo-EM maps were sharpened by using Sharpen tools in cryoSPARC and with the B factor reported in non-uniform refinement or Local Refinement. All the map figures were generated with the ChimeraX program.

### Model building, refinement and analysis

For model building, the initial model of each individual protein subunit of the TOC-TIC supercomplex was predicted by using AlphaFold2 (ref. ^45^). The predicted models were used as references or docked directly in the cryo-EM map (if they have features fitting well with the densities) during the model building process. The sharpened maps were used as references to guide the model building process, while the original maps were used for model refinement. All identifiable parts of the predicted models matching with the local density features were selected and placed in the map by fitting them as rigid bodies in the map with UCSF Chimera 1.16 (ref. ^46^) or WinCoot 0.9.4 (ref. ^47^). Subsequently, the models were manually extended, adjusted and rebuilt in WinCoot by referring to the amino acid sequences, secondary structure prediction as well as the local densities. For Tic20 and Tic214, the predicted models of their transmembrane domains were docked firstly into the corresponding region of the cryo-EM map and adjusted manually to fit each amino acid residue in the map. Afterward, the models were extended at the N-terminal and C-terminal regions according to the amino acid sequences and the map features. For Tic214, the large intermembrane-space domain has a structure largely different from the predicted model. To build its model, characteristic motifs of different regions, such as the QWWRTKQR motif in the 1233-1240 region and the RQWWW motif in the 1475-1479 region, were identified in the map. Their models were built *ab initio* and then extended at the two termini according to the sequence and map features. For Tic100, the characteristic β-jellyroll domain (BJD) was identified first in the map and the predicted model of this part was fitted into the map, adjusted and extended at both ends. For Tic56, Ctap3 and Ctap5, segments of their predicted models in the middle regions were identified in the map first and then docked in the map, adjusted and extended. The three small inner membrane proteins (Simp1, Simp2 and Simp3) were identified by referring to the predicted models of small proteins identified in the mass spectrometry analysis and manual fitting of the characteristic segments of the predicted models in the map. Once a good fit was achieved, the model was completed by extending on both ends according to the sequence and continuous local map features.

For the models of TOC proteins, Toc34 was identified by its characteristic long transmembrane helix and the model was extended on both ends. The predicted model of Toc75 was adapted into the map by adjusting the three POTRA domains individually and the β-strands in the membrane-spanning domain. Toc90 was identified by two α-helices in the intermembrane space domain. The predicted model of this part was fitted in the map, and then combined with the adapted model of the β-barrel domain which was adjusted and fitted strand-by-strand in the map. Ctap4 was identified by three characteristic α-helices in the intermembrane space region (Leu223-Leu259) and a β-barrel domain in the outer-membrane region. After the Toc34, Toc75, Toc90, Ctap4 and Chain X (unidentified due to insufficient local map resolution) were individually built in the local map, they were combined and merged with the TIC complex model. The supercomplex model was further adjusted and refined in WinCoot manually against the global map.

For model refinement, electronic Ligand Builder and Optimization Workbench (eLBOW)^48^ was used to generate restraint information for small molecule ligands. The models were refined by using phenix.real_space_refine^49^. The refined model was checked and readjusted in WinCoot manually. The model building and refinement cycle was repeated multiple times until most of the map features were interpreted by the model and the comprehensive validation statistical indicators for structural models were satisfied.

For channel pore analysis, the structural models of TOC or TIC complexes were imported to CAVER Analyst 2.0 (ref. ^50^). The TOC complex model with Toc34, Toc75, Toc 90 and part of Tic214 was used for probing the pore inside the TOC channel. Moreover, the TIC complex model with Tic20, Tic214, Simp1, Simp3 and Ctap5 was used for the search of a potential pore inside the TIC channel. Then, the channel pores were selected, exported and converted to BILD format for visualization in ChimeraX. The electrostatic potential surface representations were calculated by using ChimeraX.

### Extraction and identification of InsP6

For extraction of InsP6 (also known as phytic acid) molecules from the purified TOC-TIC supercomplex sample, 140 mg/ml biobeads (SM2, Bio rad) were added to the sample, mixed and incubated overnight at 25 ° C. The mixture was centrifuged at 17,000×g for 10 min at 25 °C. The supernatant was removed by pipetting, and 50% (v:v) glycerol was added to the pellets so that biobeads could be separated from the precipitated proteins. Subsequently, the mixture was centrifuged again, and the precipitate was retained. The precipitate was diluted by adding water to the same volume as the supernatant. 10 M NaOH and 500 mM EDTA Na were added to the suspension to final concentrations of 250 mM and 50 mM, respectively. After being dried under vacuum, the sample was dissolved in a mixture of methanol and water (1:1, v:v). The mixture was centrifuged at 17,000×g for 1 min at 25 ℃. The supernatant was collected immediately and used for LC/MS analysis. The standard sample of phytic acid (P8810, Sigma) was dissolved in water to 10 mM first and then diluted to 75 nM for LC/MS.

The sample extracted from the purified TOC-TIC supercomplex preparation or the standard sample was analyzed by using a high-performance liquid chromatograph (HPLC) (1290 Infinity, Agilent Technologies) coupled with a high-resolution mass spectrometer (6350 Accurate-Mass Q-TOF LC/MS, Agilent Technologies). The HPLC separation was carried out on a C18 column (100 mm×2.1 mm, Kinetex 2.6 μm C18 100 Å, Phenomenex) at a flow rate of 0.2 ml/min at 25 °C. The mobile phase consisted of two components, namely phase A (0.1% Formic acid aqueous) and phase B (0.1% Formic acid Acetonitrile). The sample on the column was first eluted for 13 min with a linear gradient from 1% to 70 % B and held for 4 min, followed by equilibration in 1%B. The MS analysis was performed in positive ion mode (electrospray ionization, ESI), with a scan range of 380–1500 m/z for MS, or 20–1500 m/z for MS/MS. Ion spray voltage was set at 3.5 kV in the positive mode, and the source temperature was 325 ℃, with nebulizer gas set at 35 psi and flow rate at 5 l/min. In the product ion mode, the collision energy was set at 30 V. Analysis of the LC/MS data was performed by using Peakview 2.1 software, and visualized by Matplotlib. The sample and standard had the same peak patterns of MS and MS/MS. Five main product ion peaks at m/z of 773.85, 768.90, 751.56, 398.43 and 393.47 were detected, which may correspond to the metal complex of phytic acid and its degradation products.

### Reporting summary

Further information on research design is available in the Nature Research Reporting Summary linked to this paper.

## Data availability

The cryo-EM maps of the *Chlamydomonas reinhardtii* TOC-TIC supercomplex and the TOC complex, and their corresponding atomic coordinates have been deposited in the Electron Microscopy Data Bank and the Protein Data Bank under the accession codes EMD-33528 and 7XZI, EMD-33529 and 7XZJ respectively. All data analyzed during this study are included in this Article and its Supplementary Information. Any other relevant data are available from the corresponding author upon reasonable request.

## Acknowledgements

We thank Xiao-Jun Huang, Bo-Ling Zhu, Xu-Jing Li, De-Yin Fan, Long-Long Zhang and other staff members at the Center for Biological Imaging (CBI), Core Facilities for Protein Science at the Institute of Biophysics, Chinese Academy of Science (IBP, CAS) for the support in cryo-EM data collection; Zhen-Sheng Xie, Xiang Ding, Li-li Niu at the CAS Research Platform for Protein Sciences for the support in mass spectrometry data collection and analysis. Xiao-Bo Liang and Xiu-Ying Liu for technical support on biochemistry, cell culture, sample preparation. We also thank Silvia Ramundo for sharing the HA-tagged Tic214 strain of *C. reinhardtii* and Wenqiang Yang for recovering and aliquoting the cells.

This work was funded by the Chinese Academy of Sciences Project for Young Scientists in Basic Research (YSBR-015), the National Natural Science Foundation of China (31925024), the Strategic Priority Research Program of CAS (XDB37020101 to Z.L.) and the National Key R&D Programme of China (2017YFA0503702).

## Author information

### Affiliations

National Laboratory of Biomacromolecules, CAS Center for Excellence in Biomacromolecules, Institute of Biophysics, Chinese Academy of Sciences, Beijing 100101, China; College of Life Sciences, University of Chinese Academy of Sciences, Beijing 100049, China.

Hao Liu, Anjie Li, Zhenfeng Liu

Department of Molecular Biology, University of Geneva, Geneva CH-1211, Switzerland; Department of Plant Biology, University of Geneva, Geneva CH-1211, Switzerland.

Jean-David Rochaix

### Contributions

Hao Liu expressed and purified the protein complex sample, prepared the sample for the cryo-EM study, carried out cryo-EM data collection and prepared the figures. Hao Liu and Anjie Li processed the cryo-EM data. Zhenfeng Liu and Hao Liu built and refined the atomic model, analyzed the structure. Hao Liu designed the mass spectrometry study and prepared the sample for the MS analysis. Zhenfeng Liu , Hao Liu, Anjie Li wrote the initial draft and all authors edited the manuscript. Jean-David Rochaix was involved in designing the project. Zhenfeng Liu conceived and coordinated the research project.

## Additional information

Correspondence and requests for materials should be addressed to Zhenfeng Liu.

## Extended data figures 1-10

**Extended Data Fig. 1.**
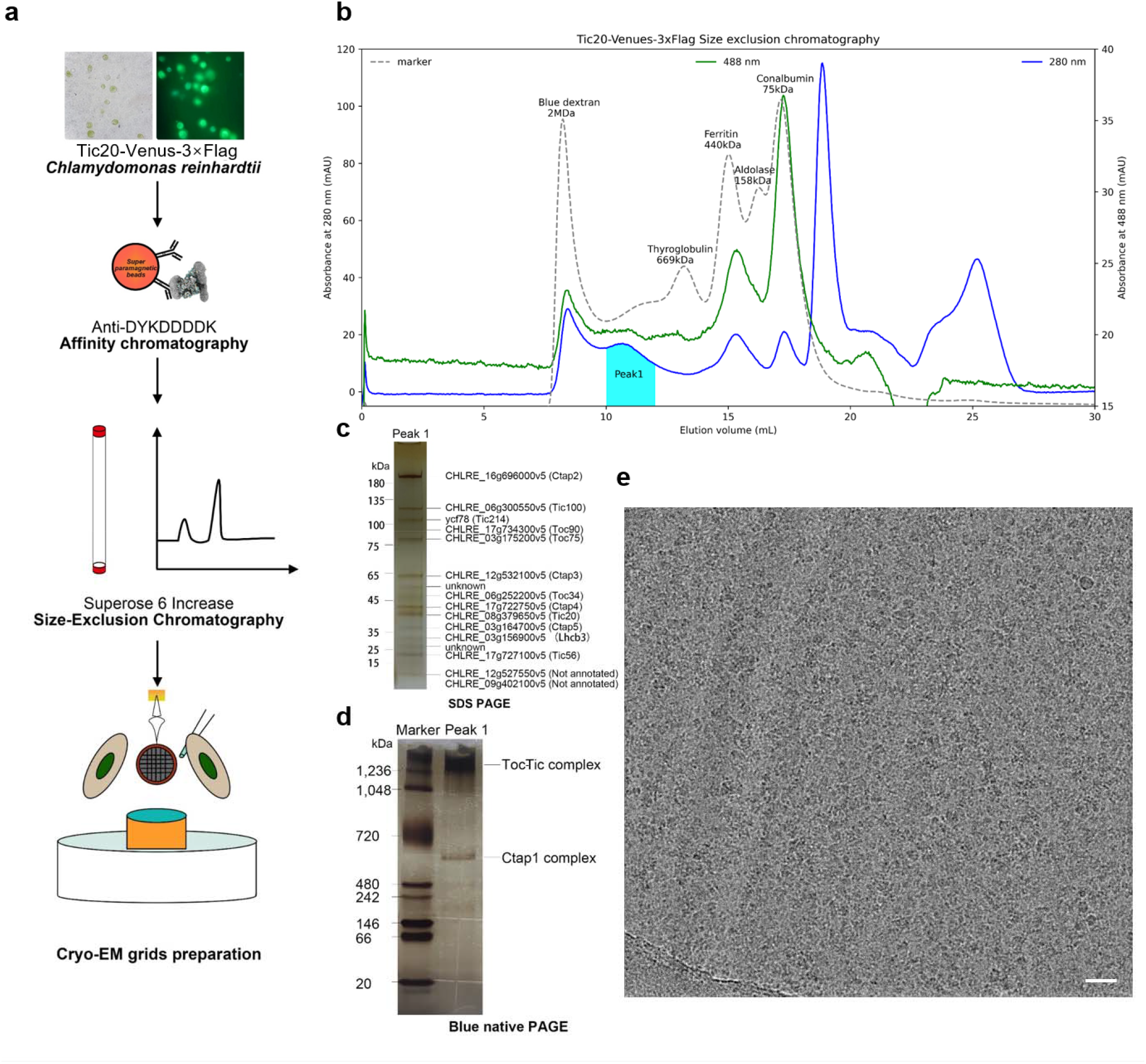
Purification and characterization of the TOC-TIC supercomplex from *C. reinhardtii*. **a**, A schematic diagram illustrating the overall flowchart of the protein purification and cryo-EM grid preparation protocol. **b**, The profiles of size exclusion chromatography loaded with the sample eluted from the Anti-DYKDDDDK beads. The peak 1 fractions marked in the cyan area were pooled, concentrated and used for cryo-EM grid preparation. The blue solid line and green solid lines indicate the absorbance measured with the UV-Vis detector at 280 nm or 488 nm (the values and units are labeled on the Y axes on left and right sides) respectively. The gray dashed line is the size exclusion chromatography of the Gel Filtration Cal Kit High Molecular Weight sample (28-4038-42, Cytiva) loaded as a reference. **c** and **d**, The SDS-PAGE (c) and blue-native PAGE (d) analyses on the peak 1 fraction from the size exclusion chromatography. Silver staining of the gel was applied for visualization of the proteins. The identities of the protein bands are labeled on the right side according to the identification results obtained through mass spectrometry on the in-gel tryptic digestion products. **e**, A representative cryo-EM micrograph of the TOC-TIC supercomplex. Scale bar, 20 nm.

**Extended Data Fig. 2.**
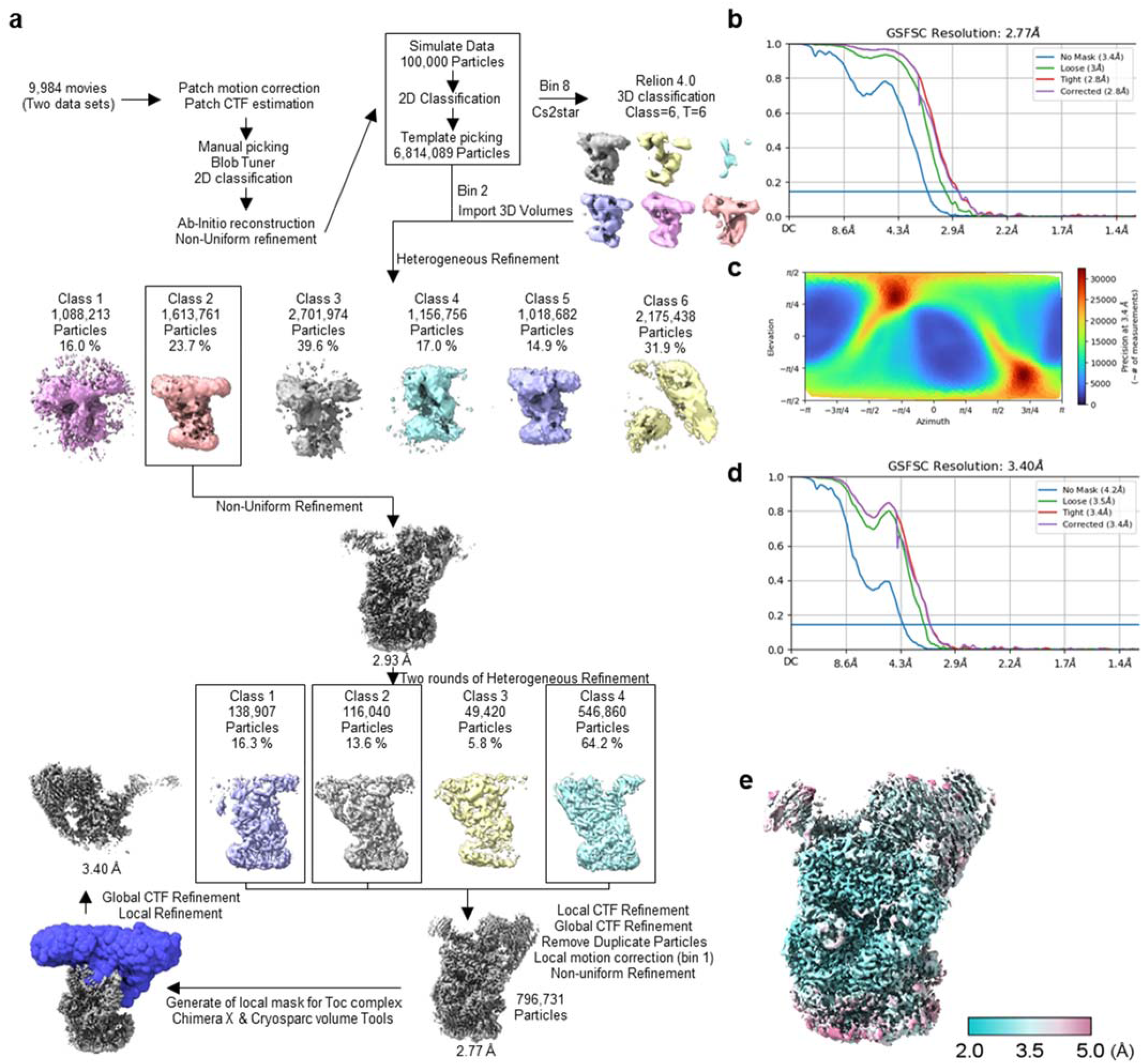
Cryo-EM data processing of the TOC-TIC supercomplex. **a**, Flowchart of the cryo-EM data processing and analysis of the TOC-TIC supercomplex. (Please refer to the ‘Cryo-EM data processing and refinement’ section in Methods for more technical details). The two datasets were collected at different spots of the same grid sample. **b** and **d** The gold-standard Fourier shell correlation (GSFSC) curves of the TOC-TIC supercomplex map (b) and the local map around the TOC complex. **c**, The posterior precision directional distribution plot of the particles of the final reconstruction generated by cryoSPARC. **e**, The local resolutions for different regions of the final overall reconstruction of the TOC-TIC supercomplex. The local resolutions were estimated by cryoSPARC and visualized in ChimeraX.

**Extended Data Fig. 3.**
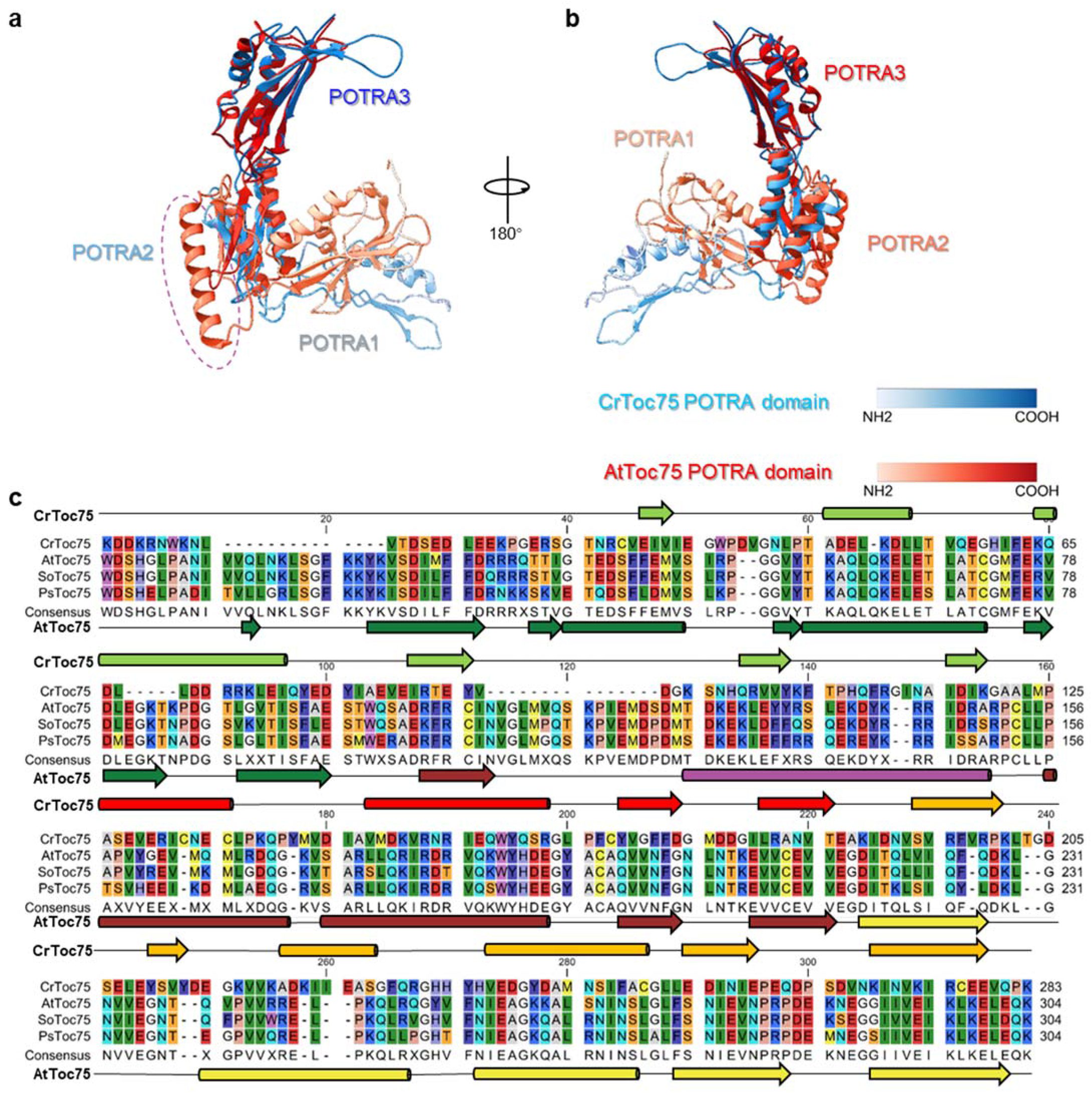
Comparison of the POTRA domains of *C. reinhardtii* Toc75 with the corresponding ones from plant Toc75. **a** and **b**, Superposition of the cryo-EM structure of *Cr*Toc75 POTRA domains with the crystal structure of *Arabidopsis* Toc75 POTRA domains (PDB: 5UAY). The superposition was generated by using Chimera X matchmaker. **c**, Alignment of the amino acid sequences of the POTRA domains of Toc75 from *Chlamydomonas reinhardtii*, *Arabidopsis thaliana*, *Spinacea oleracea* and *Pisum sativum*. The sequence alignment was carried out by using the CLC Sequence Viewer 8. Note that *At*Toc75 has an α-helix (marked in purple dotted circle in a and the purple cylinder in c) after the first β sheet of POTRA 2, which is not found in *Cr*Toc75 and has a sequence largely different from the corresponding region in *Cr*Toc75.

**Extended Data Fig. 4.**
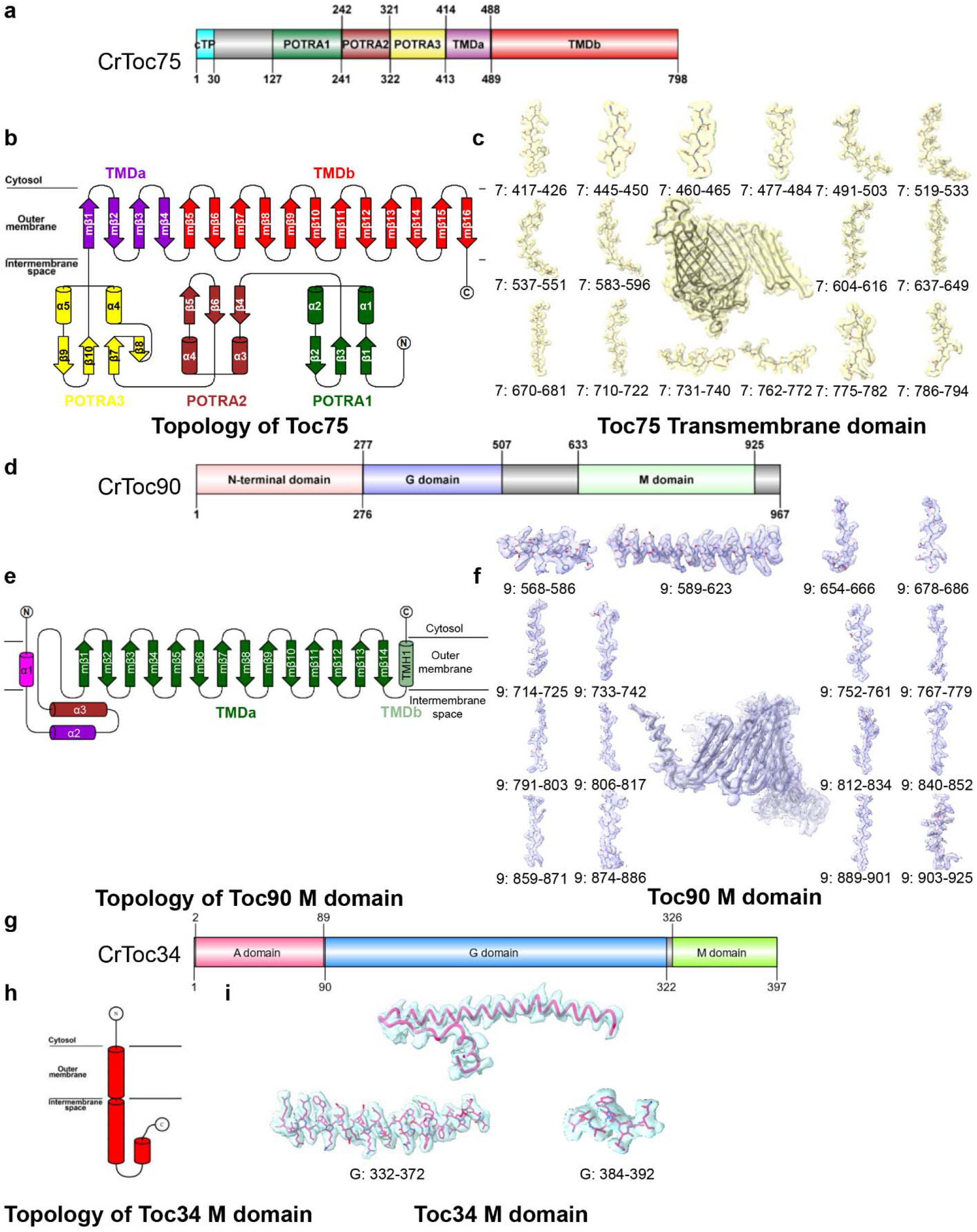
Domain organization, secondary structure topology and cryo-EM densities of Toc75 , Toc90 and Toc34. **a,** Domain organization of Toc75 protein. **b,** Topology of the secondary structures in Toc75. **c**, Cryo-EM densities of various parts in Toc75. **d**, Domain organization of Toc90 protein. **e**, Topology of the secondary structures in Toc90. **f**, Cryo-EM densities of various parts in Toc90. **g**, Domain organization of Toc34 protein. **h**, Topology of the secondary structures in Toc34. **i**, Cryo-EM densities of various parts in Toc34.

**Extended Data Figure 5.**
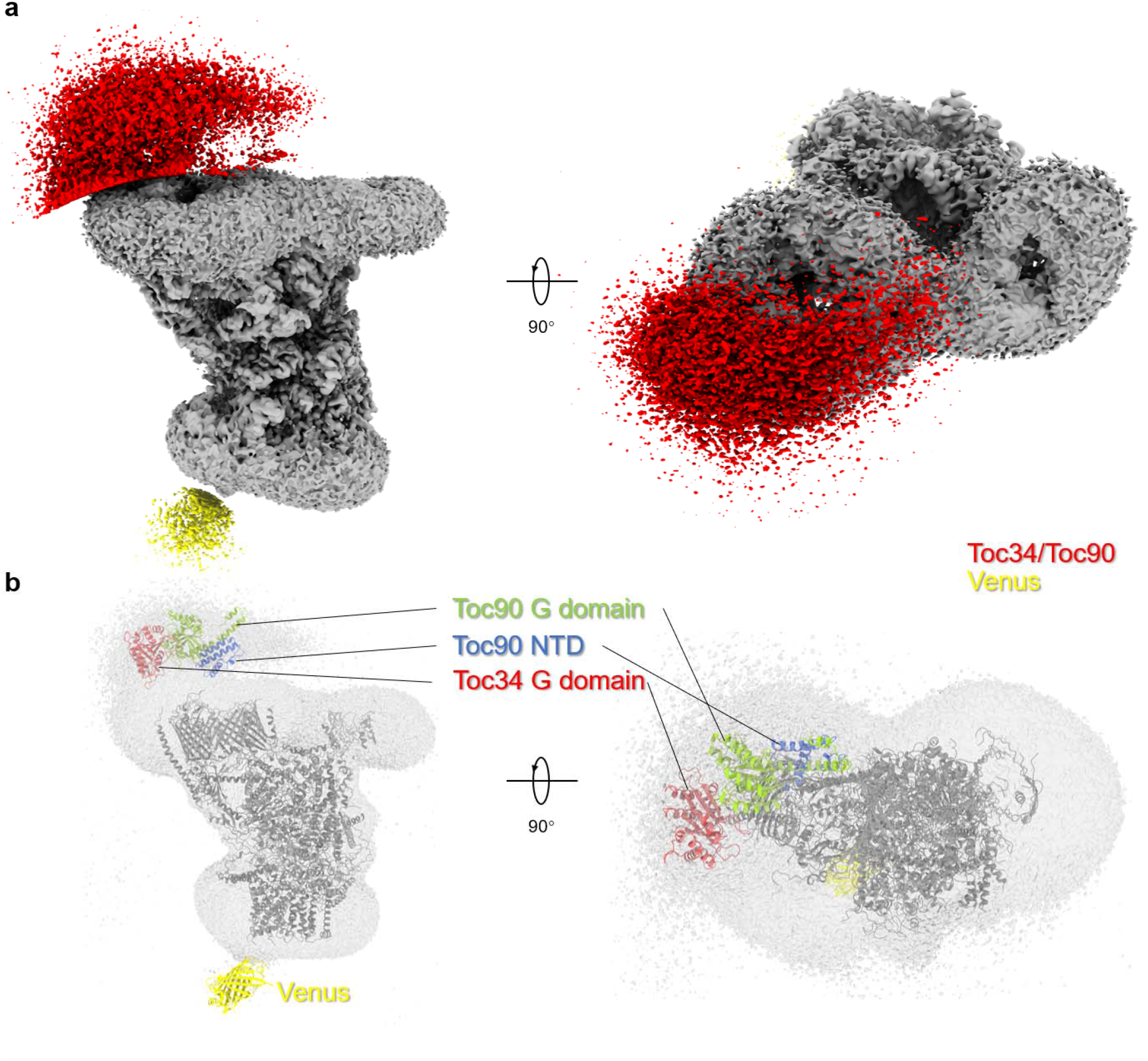
The cryo-EM densities in the cytoplasmic region above the TOC complex. **a,** The cryo-EM map is contoured at a level of 0.02 V for the cytoplasmic domain (red), 0.015 V for the stromal domain (yellow) or 0.1 V for the other regions (gray). When contoured at low levels, the weak densities on the cytosolic and stromal sides are visible. **b,** The cryo-EM map is contoured at 0.02 V level to show the weak densities on the cytosolic side. The structural models of the G domain of Toc34, and the G domain and the N-terminal domain (NTD) of Toc90 predicted by Alphafold2 are docked in the map. A putative position of the yellow fluorescence protein (Venus) fused to the C-terminal region of Tic20 is also shown. As the densities in the cytosolic and stromal regions are fragmented and too weak to serve as a reliable reference, the models of the cytosolic domains and Venus were not included in the final model of the TOC-TIC supercomplex.

**Extended Data Figure 6.**
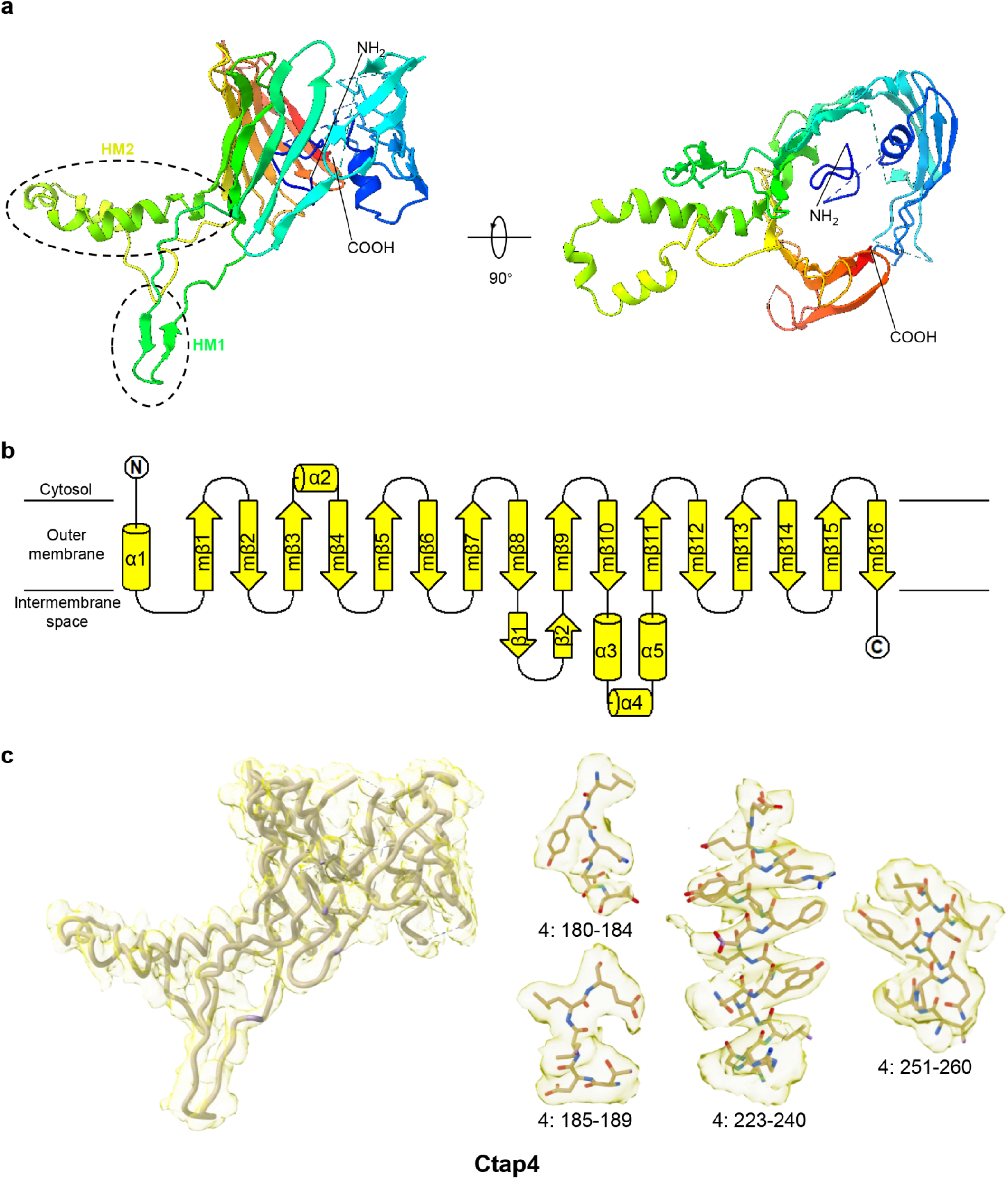
The structure, topology and cryo-EM densities for Ctap4. **a,** Side view and top view of Ctap4 structure. HM1 and HM2, hydrophilic motifs 1&2. **b,** Topological arrangement of the secondary structures in Ctap4 as predicted by Alphafold2. **c,** Cryo-EM densities of various parts in Ctap4.

**Extended Data Fig. 7.**
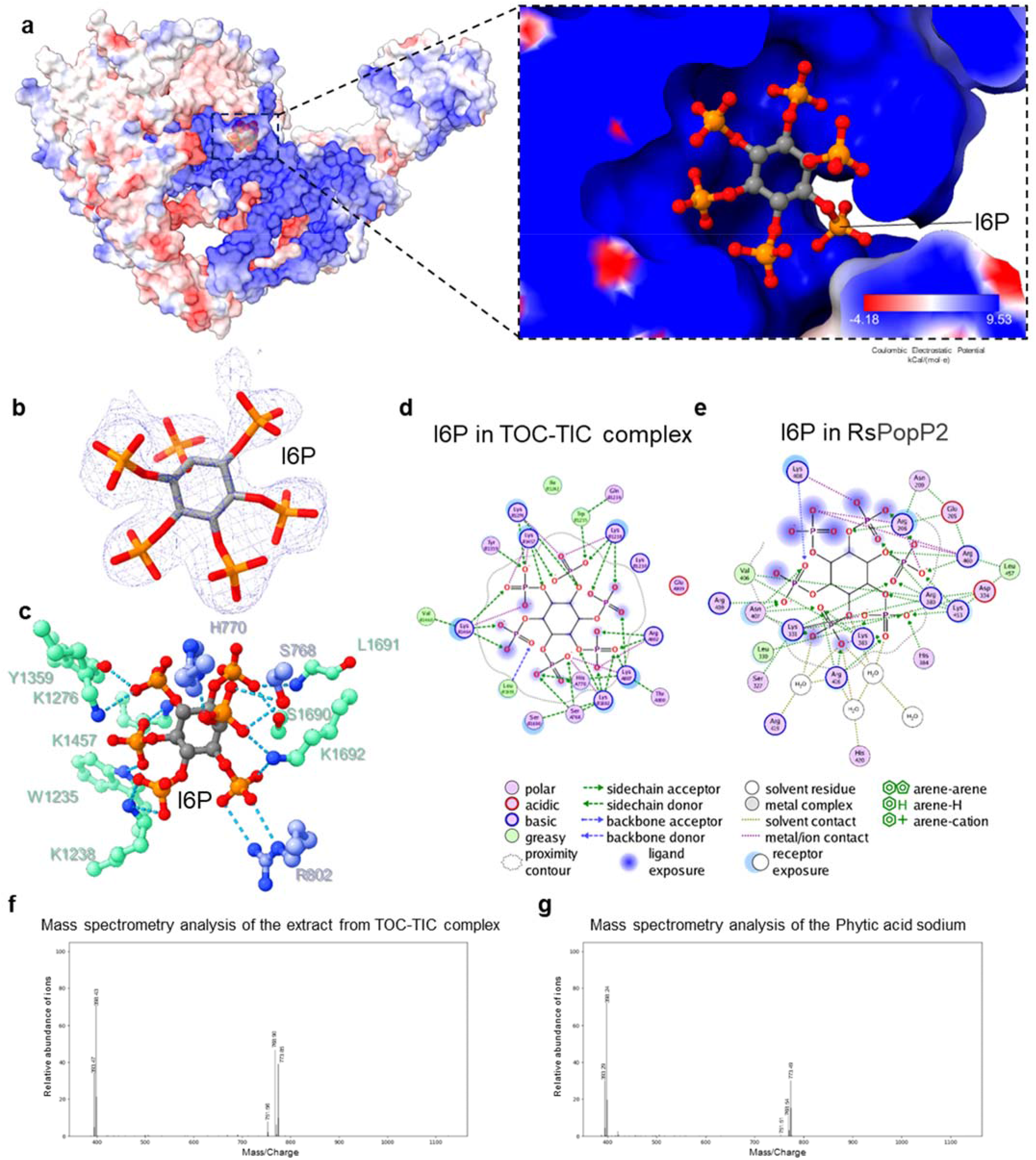
Location and identification of an InsP6 molecule in the TOC-TIC supercomplex from *C. reinhardtii*. **a,** Electrostatic surface representation of the TOC complex hosting an InsP6-binding site. The zoom-in view on the right side shows an InsP6 molecule accommodated in a positive charge-rich cavity. The electropositive surface of the cavity is complementary to the negative charges carried by the phosphate groups of InsP6. **b,** The cryo-EM density of the InsP6 molecule with the stick model superposed. The cryo-EM map is contoured at 0.25 V. **c,** Interactions of InsP6 with adjacent amino acid residues from Tic214 (green) and Toc90 (blue) in the TOC-TIC supercomplex. **d and e,** The 2D interaction map of InsP6 in the TOC-TIC supercomplex (d) or *Rs*PopP2 (PDB:5W3X, e) calculated by using the MOE program. *Rs*PopP2 is a bacterial effector protein produced by the plant pathogen *Ralstonia solanacearum* and hosts an InsP6-binding site similar to the one found in the TOC complex. The phosphate groups of InsP6 molecules are surrounded by positively-charged residues in both cases. **f** and **g,** Mass spectrometry analysis of the extract from the TOC-TIC supercomplex sample (f) or the standard sample of phytic acid (sodium salt, g).

**Extended Data Fig. 8.**
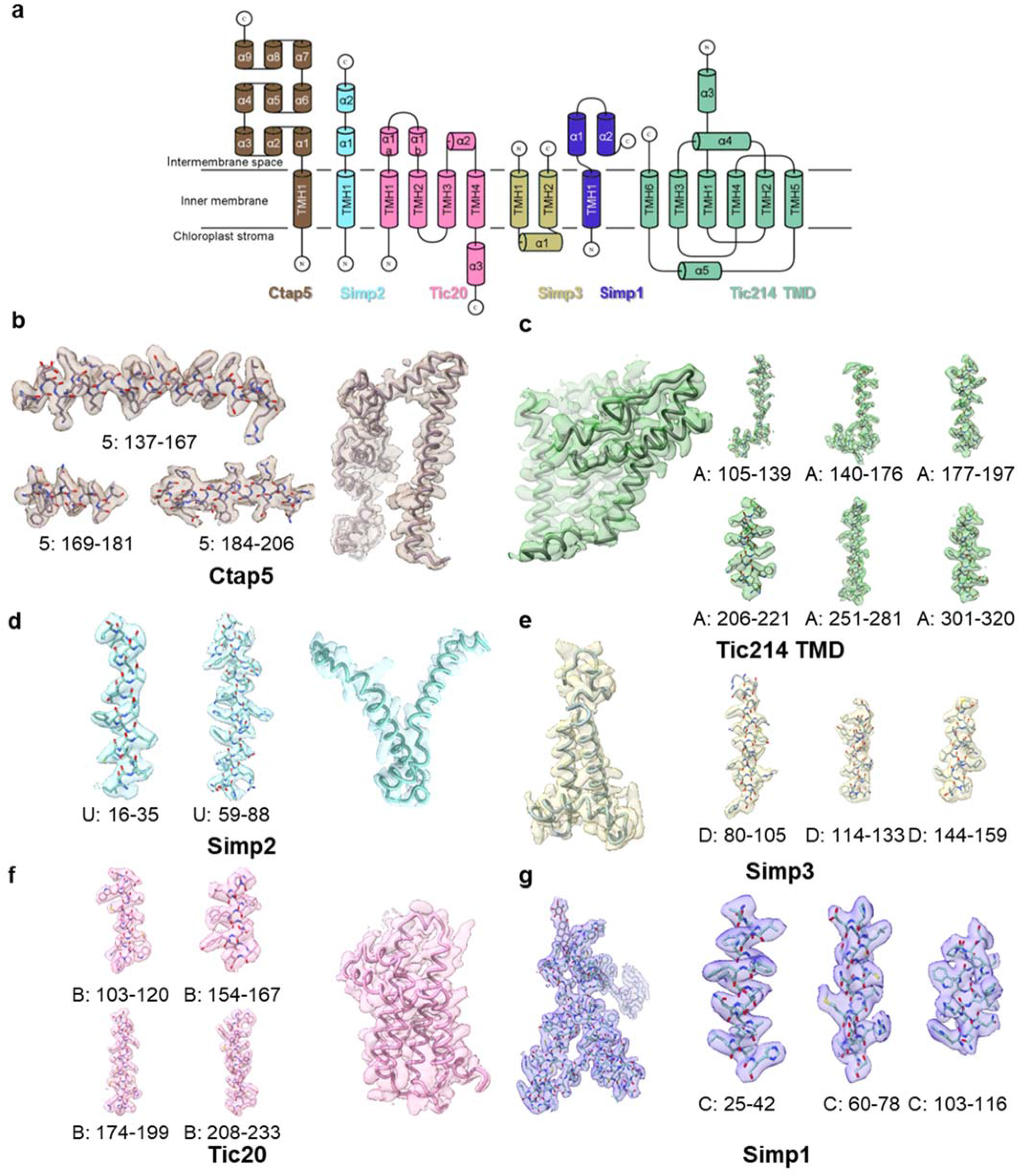
Topology and cryo-EM densities of the six proteins involved in forming the TIC complex. **a,** Topological arrangement of secondary structures in Ctap5, Simp2, Tic214, Tic20, Simp3 and Simp1 proteins. **b-g,** Cryo-EM densities of various parts in the six proteins.

**Extended Data Fig. 9.**
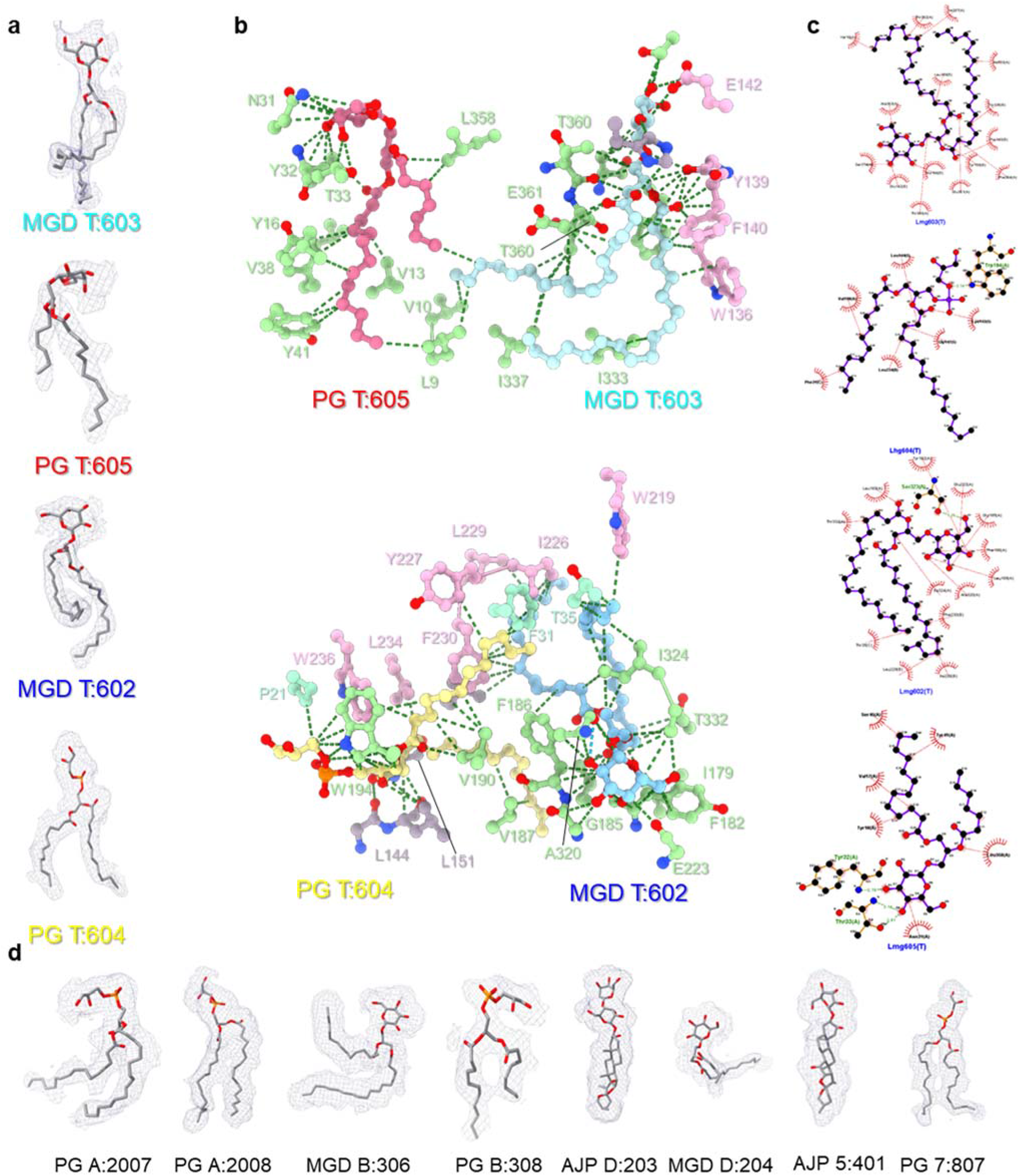
The lipid and detergent molecules associated with the TOC-TIC supercomplex. **a,** The cryo-EM densities of four central lipid molecules located in or above the putative pore of the TIC complex. **b,** Interactions of the four lipid molecules with adjacent amino acid residues. Green, Tic214; Pink, Tic20; brown, Ctap5. **c**, LigPlot analyses showing the hydrogen bonds and hydrophobic interactions formed between the four lipid molecules and nearby amino acid residues. **d**, The cryo-EM densities of other peripheral lipid molecules associated with Tic214(A), Tic20(B), Simp3(D), Ctap5(5) and Toc75(7).

**Extended Data Fig. 10.**
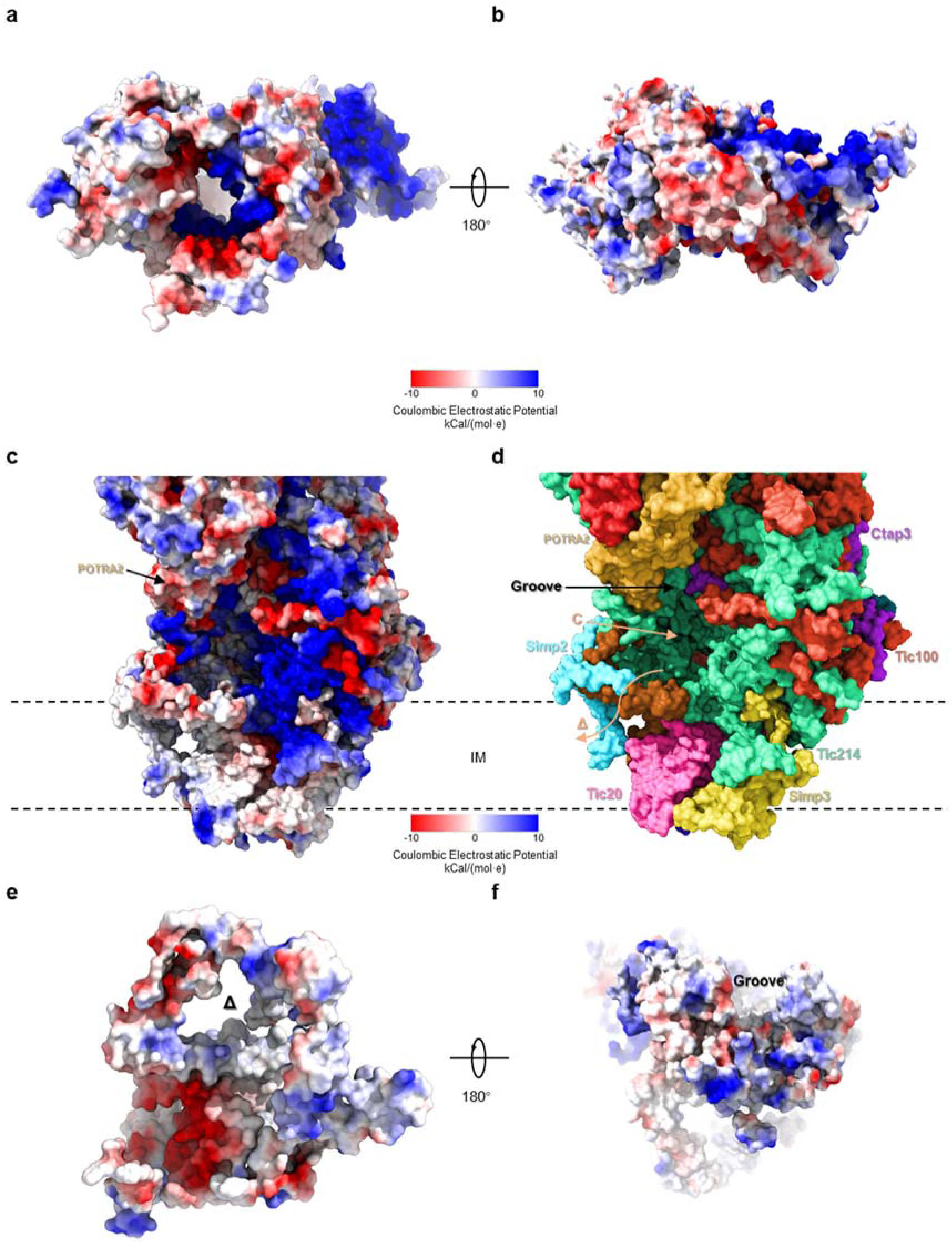
The surface presentations of the TOC complex, the intermembrane space domain connecting the TOC channel with the TIC channel, and the TIC complex. **a and b,** The electrostatic potential surface of the TOC complex viewed from the cytoplasmic and intermembrane space sides, respectively. **c,** The electrostatic potential surface of the ISC connecting the TOC complex with the TIC complex. **d,** The presence of a surface groove connecting the channels of the TOC and TIC complexes. The local regions labeled by “C” and “Δ” are the cavity above the central funnel of the TIC complex and the triangular entrance formed by Ctap5, Simp2 and Tic214, respectively. **e and f,** The electrostatic potential surface of the TIC complex viewed from the intermembrane space and stromal sides, respectively.

