## Supplementary Tables 1 and 2 for "Architecture of chloroplast TOC-TIC translocon supercomplex"

**Supplementary Table 1 A summary of each individual protein subunit identified in the TOC-TIC supercomplex from *C. reinhardtii*.**

| **Gene name** | **Entry** | **Protein name** | **Chain ID** | **Length (aa)** | **SDS PAGE PMF score** | **BN PAGE unique peptides** |
| --- | --- | --- | --- | --- | --- | --- |
| CHLRE_06g300550v5 | A0A2K3DQY7 | Tic100 | E | 955 | 138 | 24 |
| ycf78 | P36495 | Tic214 | A | 1995 | 36 | 77 |
| CHLRE_17g734300v5 | A0A2K3CR90 | Toc90 | 9 | 967 | 108 | 19 |
| CHLRE_03g175200v5 | A8IE32 | Toc75 | 7 | 798 | 212 | 22 |
| CHLRE_12g532100v5 | A0A2K3D4W3 | Ctap3 | 3 | 462 | 139 | 14 |
| CHLRE_06g252200v5 | A8HYJ1; A0A2K3DM34 | Toc34 | G | 397 | 74 | 2 |
| CHLRE_17g722750v5 | A8J6H7 | Ctap4 | 4 | 363 | 120 | 8 |
| CHLRE_03g164700v5 | A0A2K3DWN5; A8IFJ3 | Ctap5 | 5 | 383 | 49 | 11 |
| CHLRE_17g727100v5 | A8J6R5 | Tic56 | F | 244 | 95 | 8 |
| CHLRE_12g527550v5 | A8J5D4 | Simp2 | U | 124 | - | 4 |
| CHLRE_09g402100v5 | A8J1J3 | Simp1 | C | 127 | - | 3 |
| CHLRE_02g080250v5 | A0A2K3E0K3 | Simp3 | D | 187 | - | 2 |
| CHLRE_08g379650v5 | A8IZ79 | Tic20 | B | 259 | - | 2 |

The proteins in the purified TOC-TIC supercomplex sample were subject to mass spectrometry (MS) analysis. The MALDI-TOF/TOF was used for the tryptic digestion products of individual protein bands excised from the SDS-PAGE gel, and the nano LC-Q EXACTIVE was used for the tryptic digestion products of the supercomplex band from BN-PAGE gel. The proteins were identified by considering the peptide mass fingerprinting (PMF) score, unique peptides found in the BN-PAGE band and fitting of the models in the cryo-EM densities.

**Supplementary Table 2 Cryo-EM data collection, refinement and validation statistics.**

|  | #1 TOC-TIC supercomplex  (EMDB-33528)  (PDB 7XZI) | #2 TOC complex (local)  (EMDB-33529)  (PDB 7XZJ) |
| --- | --- | --- |
| **Data collection and processing** |  |  |
| EM equipment | FEI Titan Krios G3 | |
| Detector | Gatan K2 | |
| Magnification | 105,000× | 105,000× |
| Voltage (kV) | 300 | 300 |
| Electron exposure (e–/Å^2^) | 50 | 50 |
| Number of frames/movie | 33 | 33 |
| Defocus range (μm) | -1.5~1.8 | -1.5~1.8 |
| Pixel size (Å) | 0.675 | 0.675 |
| **Reconstruction** |  |  |
| Software | Cryosparc v3.3 & Relion 4.0 | |
| Symmetry imposed | C1 | C1 |
| Initial particle images (no.) | 6,814,089 | 6,814,089 |
| Final particle images (no.) | 796,731 | 796,731 |
| Map resolution (Å)  FSC threshold | 2.77  0.143 | 3.40  0.143 |
| **Model building** |  |  |
| Software | WinCoot 0.9.4 & AlphaFold2 | |
| **Refinement** |  |  |
| Software | PHENIX 1.20.1-4487 | |
| Model resolution (Å)  FSC threshold | 3.3  0.5 | 3.8  0.5 |
| Map sharpening *B* factor (Å^2^) | -85.6 | -96.0 |
| Model composition  Non-hydrogen atoms  Protein residues  Lipids/Detergents/other ligand | 39,308  4,838  16/6/1 | 15,434  1,899  5/4/1 |
| *B* factors (Å^2^)  Protein  Ligand | 115.80  84.66 | 72.56  42.47 |
| R.m.s. deviations  Bond lengths (Å)  Bond angles (°) | 0.009  1.435 | 0.008  1.355 |
| Validation  MolProbity score  Clashscore  Poor rotamers (%) | 3.25  20.24  15.88 | 3.09  24.22  5.09 |
| Ramachandran plot  Favored (%)  Allowed (%)  Disallowed (%) | 90.73  8.22  1.05 | 84.15  12.39  3.46 |
